# Integrative multi-dimensional characterization of striatal projection neuron heterogeneity in adult brain

**DOI:** 10.1101/2023.05.04.539488

**Authors:** Jenesis Gayden, Stephanie Puig, Chaitanya Srinivasan, BaDoi N. Phan, Ghada Abdelhady, Silas A. Buck, Mackenzie C. Gamble, Hugo A. Tejeda, Yan Dong, Andreas R. Pfenning, Ryan W. Logan, Zachary Freyberg

**Affiliations:** Center for Neuroscience, University of Pittsburgh, Pittsburgh, PA 15213, USA; Department of Psychiatry, University of Pittsburgh, Pittsburgh, PA 15213, USA; Department of Psychiatry, University of Massachusetts Chan Medical School, Worcester, MA 01605, USA; Department of Computational Biology, Carnegie Mellon University, Pittsburgh, PA 15213, USA; Medical-Scientist Training Program, University of Pittsburgh, Pittsburgh, PA 15213, USA; Department of Pharmacology and Experimental Therapeutics, Boston University School of Medicine, Boston, MA, USA; Molecular and Translational Medicine, Department of Medicine, Boston University School of Medicine, Boston, MA, USA; Unit on Neuromodulation and Synaptic Integration, National Institute of Mental Health, Bethesda, MD 20894, USA; Department of Neuroscience, University of Pittsburgh, Pittsburgh, PA 15260, USA; Neuroscience Institute, Carnegie Mellon University, Pittsburgh, PA 15213, USA; Department of Neurobiology, University of Massachusetts Chan Medical School, Worcester, MA 01605, USA; Department of Cell Biology, University of Pittsburgh, Pittsburgh, PA 15213, USA

**Author notes:** Corresponding author: Zachary Freyberg, M.D., Ph.D., 3811 O’Hara Street, BST W1640, Pittsburgh, PA 15213.

**Keywords:** Dopamine, striatum, RNAscope, RNAseq, transcriptomics

## Abstract

Striatal projection neurons (SPNs) are traditionally segregated into two subpopulations expressing dopamine (DA) D_1_-like or D_2_-like receptors. However, this dichotomy is challenged by recent evidence. Functional and expression studies raise important questions: do SPNs co-express different DA receptors, and do these differences reflect unique striatal spatial distributions and expression profiles? Using RNAscope in mouse striatum, we report heterogenous SPN subpopulations distributed across dorsal-ventral and rostral-caudal axes. SPN subpopulations co-express multiple DA receptors, including D_1_ and D_2_ (D1/2R) and D_1_ and D_3_. Our integrative approach using single-nuclei multi-omics analyses provides a simple consensus to describe SPNs across diverse datasets, connecting it to complementary spatial mapping. Combining RNAscope and multi-omics shows D1/2R SPNs further separate into distinct subtypes according to spatial organization and conserved marker genes. Each SPN cell type contributes uniquely to genetic risk for neuropsychiatric diseases. Our results bridge anatomy and transcriptomics to offer new understandings of striatal neuron heterogeneity.

## Introduction

Striatal dopamine (DA) is important for motivation, reward, and motor control^1–3^. Dysfunctions in DA signaling in striatum are strongly associated with numerous prevalent neurologic and psychiatric diseases including schizophrenia, substance use disorder, mood disorders, and Parkinson’s disease^4–11^. Striatal DA signaling is mediated by DA receptors expressed by GABAergic spiny projection neurons (SPNs). These DA receptors are categorized into 2 major classes: Gα_s_-coupled D_1_-like G protein-coupled receptors (GPCRs) comprising D_1_ and D_5_ receptors (D1R, D5R) and Gα_i_-coupled D_2_-like GPCRs including D_2_, D_3_, and D_4_ receptors (D2R, D3R, D4R)^12^.

Traditionally, SPNs have been segregated into two subpopulations based on expression of D_1_-like or D_2_-like receptors^12–14^. Indeed, SPNs expressing D_1_-like receptors differ from SPNs that express D_2_-like receptors at cellular, circuit, and behavioral levels^15–21^. However, many questions remain concerning this dichotomous separation. For example, does expression of specific D_1_-like versus D_2_-like DA receptors underlie the differences between SPN subpopulations? Increasing evidence also suggests that the dichotomous division of SPNs according to their expression of D_1_-like or D_2_-like receptors is not clear-cut^22,23^. Instead, SPNs may be more heterogenous than previously appreciated. Earlier findings have suggested the existence of additional striatal SPN subpopulations apart from the traditionally defined SPNs that solely express either D1R or D2R. Consistent with this, recent work found evidence of SPNs that co-express both receptors^23–26^. Yet, it remains unclear whether these co-expressing SPNs exhibit striatal distribution patterns distinct from those of SPNs that solely express single DA receptors.

Compared to D1R and D2R, considerably less is known about the biology and anatomic distribution of striatal D3R. Functionally, striatal D3R signaling is implicated in reward, as well as in diseases including Parkinson’s disease and impulse control disorders^27–31^. Anatomically, D3R expression is more restricted than D1R and D2R, but is more highly expressed in the ventral striatum^32,33^. Similar to D1R and D2R, heterologous overexpression studies have suggested that D3R also interacts with other DA receptors^34–37^. Thus, understanding the expression patterns of D1R, D2R, and D3R may therefore shed new light on the relevance of singly- and co-expressed DA receptors within striatal circuits in health and disease.

Historically, attempts at disentangling different striatal cell type subpopulations according to DA receptor expression have been limited by the paucity of sufficiently sensitive and specific tools. Relative lack of DA receptor-specific antibodies, coupled with spatial resolution below the single-cell level using earlier methods such as autoradiography or fluorescent ligand binding^38–44^, have prevented accurate identification and mapping of distinct striatal subpopulations. Recent development of much more sensitive, quantitative approaches with single-cell resolution such as multiplex RNAscope, a form of fluorescent *in situ* mRNA hybridization, has been instrumental in overcoming these limitations. In parallel, unbiased single-cell transcriptomics and epigenomics have introduced a complementary approach to studying cellular heterogeneity in the brain, including the striatum^45–54^. While such work has provided invaluable data on striatal composition, it has also introduced a plethora of different definitions for SPNs with limited agreement^55^. Such lack of consistency in classifying SPNs has created a barrier to further investigation of distinct SPNs. This may also be due in part to limited integration of spatial distributions of the different SPN subpopulations. Therefore, to tackle these challenges, we have created a new integrative approach that combines single-nuclei multi-omic strategies across several species with RNAscope mapping in striatum. Our approach enables us to provide a simple consensus that describes SPNs across diverse datasets, and that connects these data to complementary three-dimensional spatial mapping.

Here, we establish a definitive multi-dimensional map of D1R, D2R, and D3R mRNA expression in the striatum. We also confirm DA receptor co-expressing SPN subpopulations (D1/2R, D1/3R, D2/3R) that are discrete from SPNs that solely express D1R, D2R, or D3R. Importantly, we reveal that these subpopulations have distinct spatial distributions, not only in the dorsal-ventral direction, but also along the rostral-caudal axis. These differences in distribution point to a three-dimensional (3D) gradient of DA receptor expression in adult brain. Concurrently, we combine the multi-omic single-cell datasets from the adult striatum to create a straightforward and uniform classification of the SPN cell types found in mice, rats, rhesus macaques, and humans. Lastly, we compare the epigenomic profiles of different striatal cell types with human genetic risk variants for neuropsychiatric traits to reveal these cells’ distinct associations with markers of neurological and psychiatric disorders. Overall, our results bridge anatomy and transcriptomics to offer new understandings of striatal neuron heterogeneity in health and disease.

## Results

### Establishing RNAscope mapping of mouse striatum

We comprehensively examined the mRNA expression of D1R, D2R, and D3R across entire striatal brain slices of adult mouse via multiplex RNAscope. Our choice of these DA receptors was guided by their relative abundance within the striatum in tissue expression databases of both humans and mice^56,57^. We quantified expression of D1R, D2R, and D3R in key striatal structures: the caudate putamen (CPu), the NAc core (Core), the NAc shell (Shell), and the olfactory tubercle (OT) (Figure 1). Moreover, we confirmed the specificity of the RNAscope approach in mouse striatum by testing pre-validated positive and negative controls (Figure S1).

**Figure 1.**
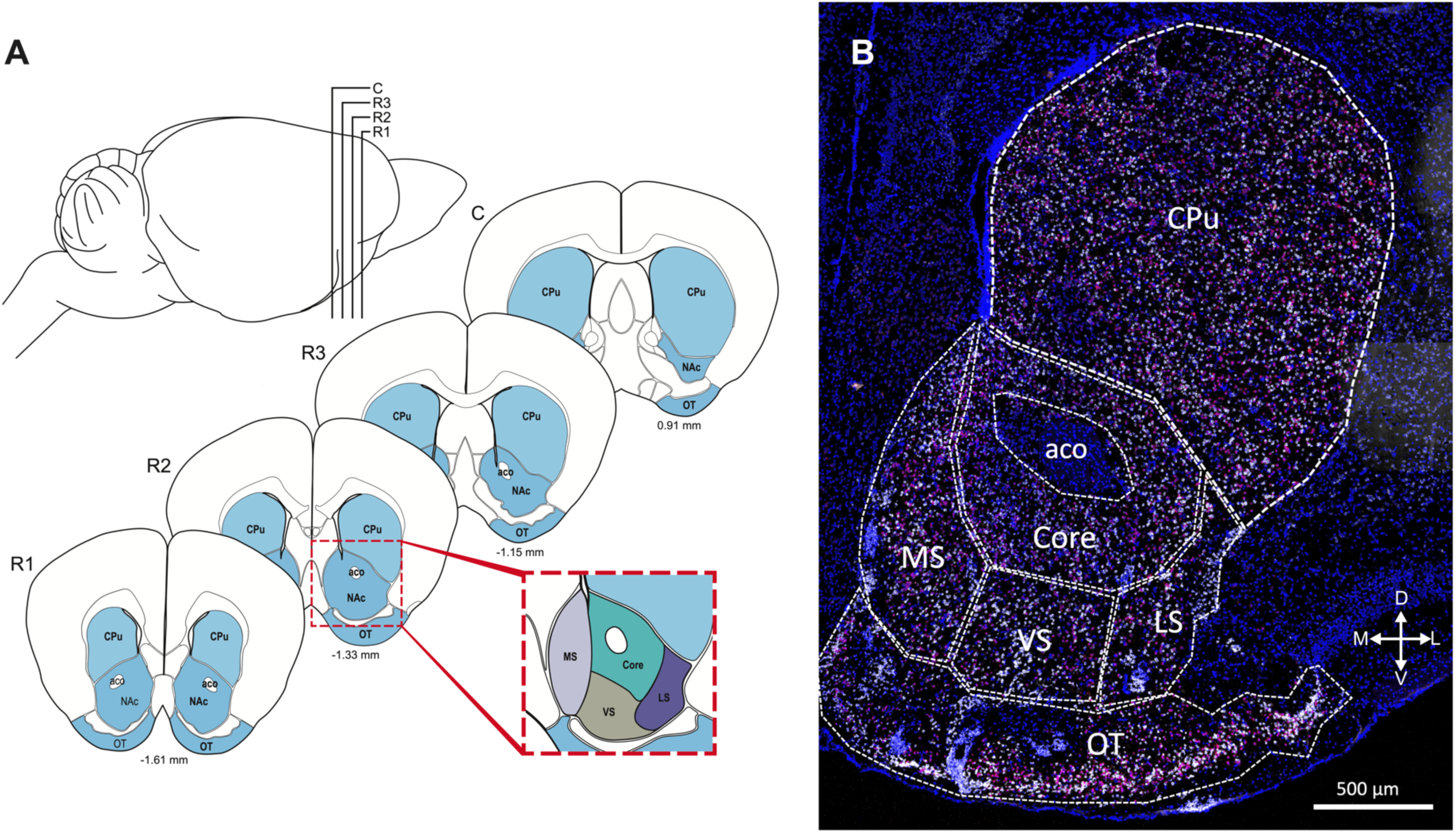
Diagrams representing the brain regions analyzed. (**A**) Coronal sections of mouse striatum were taken at Bregma -1.61 mm, referred to in the text and data as Rostral 1 (R1); -1.33 mm, Rostral 2 (R2); -1.15 mm, Rostral 3 (R3), and 0.91 mm, Caudal (C). The respective striatal subregions are annotated as follows: olfactory tubercle (OT), nucleus accumbens (NAc), anterior commissure (aco), as well as the caudate nucleus and putamen (CPu). The inset demonstrates the NAc subregions that were analyzed: NAc medial shell (MS, in purple), lateral shell (LS, in violet), ventral shell (VS, in beige), and Core (in aqua) (see Supplementary Figs. S2, S3). (**B**) Annotation of a representative RNAscope image of mouse striatum displaying the striatal regions analyzed. Scale bar = 500 μm. D=dorsal, V=ventral, L=lateral, M=medial.

### Spatial distribution of striatal dopamine D_1_, D_2_, D_3_ receptor expression in singly-expressing spiny projection neurons

We confirmed expression of D1R, D2R, and D3R throughout the striatum along both dorsal-ventral and rostral-caudal axes. We identified discrete populations of SPNs that either express a single DA receptor subtype (D1R-only, D2R-only, or D3R-only) (Figure 2, Figure S2) versus SPNs that co-express multiple receptor subtypes (D1/2R, D1/3R, D2/3R) (Figure 3, Figure S3). Focusing on SPNs expressing only one DA receptor subtype (Figure 2), there was a significant interaction between cell type, striatal subregion, and position along the rostral-caudal axis [F(17,790)=1.68, p=0.04]. When we assayed for potential sex differences in cell densities of all SPN subpopulations across the major striatal structures (CPu, NAc Core and Shell, OT) we found no significant sex differences (p>0.05). Therefore, we combined both sexes in subsequent analyses.

**Figure 2.**
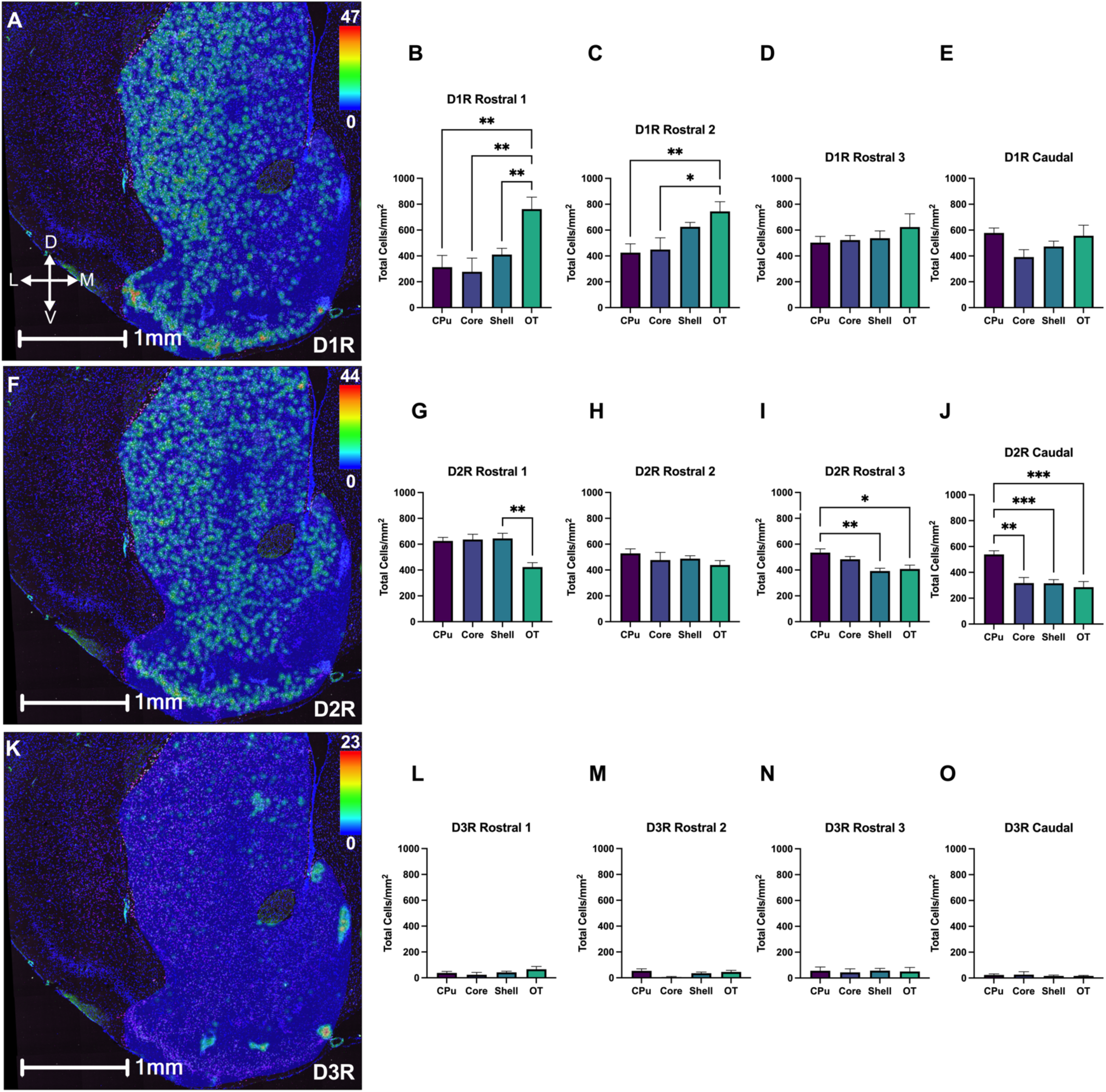
Spatial distribution of striatal dopamine D_1_, D_2_, and D_3_ receptor expression in singly expressing neurons. Analyses of multiplex RNAscope data demonstrating DA D_1_, D_2_, and D_3_ receptor expression in key striatal structures (CPu, Core, Shell, OT) along dorsal-ventral and rostral-caudal axes. (**A)** Representative heat map of the density of cells positive for D_1_ receptor (D1R) expression in singly-expressing neurons, showing robust D1R expression throughout the striatum, particularly in the OT. (**B-E**) Quantification revealed a progressive decrease in OT enrichment of D1R-only cells in a rostral to caudal direction through the striatum. (**F**) Representative heat map showing a robust distribution of cells expressing DA D_2_ receptor (D2R) mRNA throughout the striatum along the dorsal-ventral axis. **(G-J**) Quantitative analysis showed a steady decrease in the density of D2R-only neurons in the Shell and OT moving caudally (*p<0.05, **p<0.01, ***p<0.001), while remaining stable in the Core and CPu (p>0.05). (**K)** Representative heat map of SPNs expressing DA D_3_ receptor (D3R) expression in striatal neurons. D3R expression was limited, aside from enrichment in selected subregions of the Shell and OT including within the Islands of Calleja (IC). (**L-O**) Quantitative analysis revealed uniformly low D3R expression in both dorsal-ventral and rostral-caudal directions (p>0.05). For **Panels A, F, and K**, scale bar = 1 mm; heat map legends represent the number of mRNA grains within a given cell. Mean ± SEM, n=6 mice for all conditions. *p<0.05, **p<0.01, ***p<0.001. D=dorsal, V=ventral, L=lateral, M=medial.

**Figure 3.**
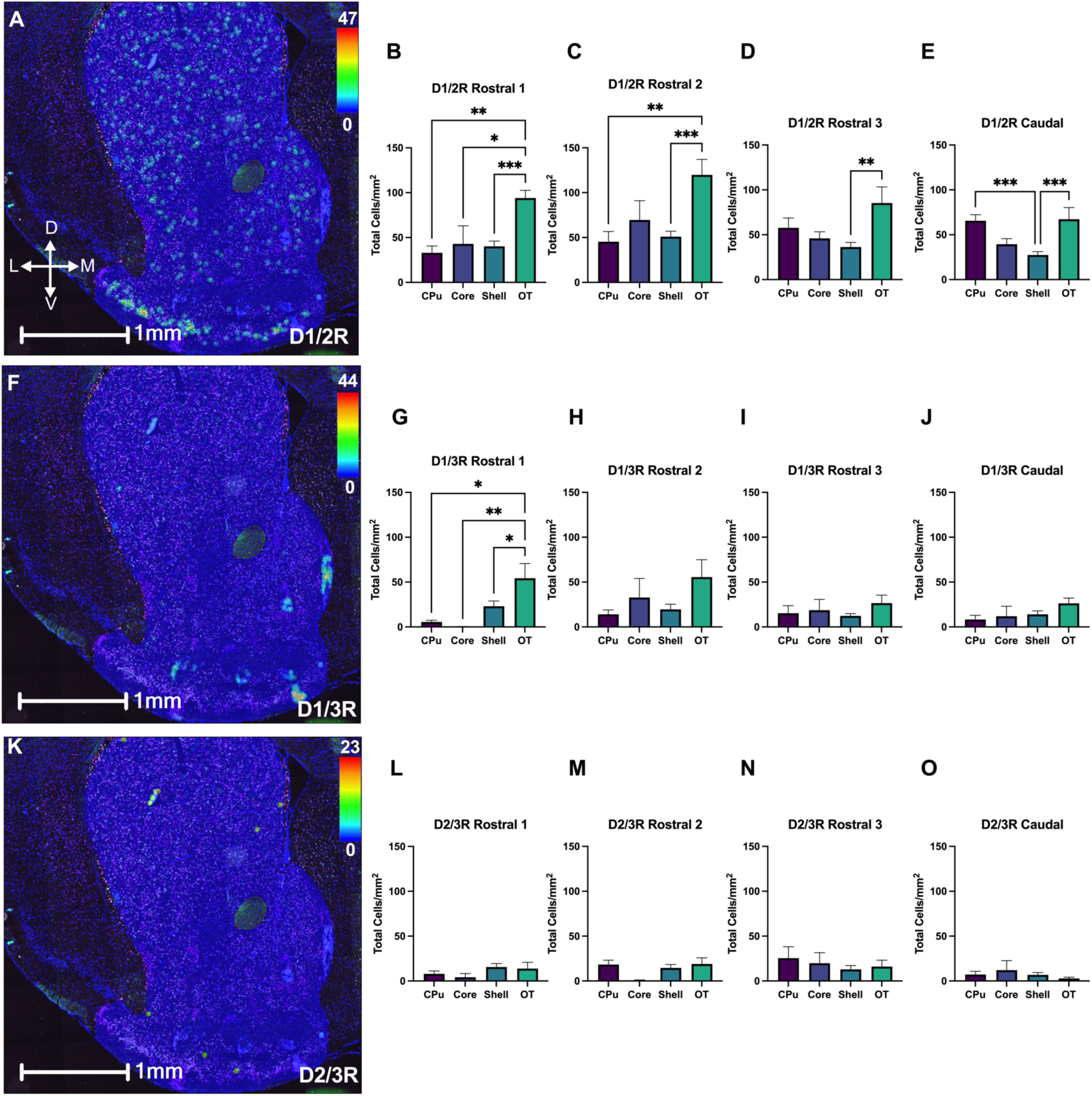
Spatial distribution of striatal dopamine D_1_, D_2_, or D_3_ receptor expression in co-expressing neurons. **(A)** Representative heat map of SPNs co-expressing D1R/D2R (D1/2R). **(B-E)** Quantitation revealed that the highest density of D1/2R SPNs is in the OT. **(F)** Representative heat map showing the density of SPNs co-expressing D1R/D3R (D1/3R). **(G-J)** Quantitative analysis demonstrated preferential D1/3R co-expression in OT neurons that diminishes moving caudally (*p<0.05, **p<0.01). (**K**) Representative heat map of SPNs co-expressing D2R/D3R (D2/3R) revealed limited pockets of co-expression in the NAc and OT. **(L-O)** Quantitative analysis found low overall levels of D2/3R SPNs with no changes in distribution along dorsal-to-ventral or rostral-to-caudal axes (p>0.05). For **Panels A, F, and K**, scale bar = 1 mm; heat map legends represent the number of mRNA grains within a given cell. Mean ± SEM, n=6 mice for all conditions. *p<0.05, **p<0.01, ***p<0.001. D=dorsal, V=ventral, L=lateral, M=medial.

#### D1R-only SPNs

D1R was expressed throughout the striatum, particularly in SPNs that solely express this receptor (D1R-only; Figure 2A). We additionally discovered distinct patterns of D1R expression across both dorsal-ventral and rostral-caudal axes. Beginning in the most rostral section (Rostral 1), the density of D1R-only cells was greatest in the OT (Figure 2B) compared to the other striatal subregions [F(3,58) = 6.24, p=0.001]. Closer analysis of the individual subsections of the NAc shell (*e.g*., medial, ventral, and lateral shell subsections; Figure 1 inset) revealed that the differences in D1R-only cell density between the OT and Shell in the Rostral 1 section were primarily driven by the lateral NAc shell (p=0.016; Supplementary Figure S2A). Moving caudally, significant differences in D1R-only cell density between the OT and other striatal subregions gradually diminished and ultimately disappeared in the Rostral 3 and Caudal sections (p>0.05; Figure 2C-2E).

#### D2R-only SPNs

D2R was widely expressed in the striatum and, similar to D1R-only cells, we identified SPNs that only expressed D2R (D2R-only; Figure 2F). In the Rostral 1 section, we observed an overall significant difference in the distribution of D2R-only cells across the striatal subregions ([F(3,56) = 4.25, p=0.009]; Figures 2F-J). There was a significantly lower density of D2R-only cells in the OT compared to the Shell (p=0.005; Figure 2G). This was due to differences in D2R-only cell density specific to the medial NAc shell (p<0.0001; Figure S2E). Interestingly, we found trends in cell distribution opposite to those of D1R-only SPNs. Specifically, moving caudally, D2R-only cell density decreased in the OT [F(3,41) = 3.88, p=0.02], NAc shell [F(3,133) = 24.23, p=<0.0001], and Core [F(3,37) = 8.09, p=0.0003). In contrast, CPu D2R-only cell density increased from Rostral 1 to Rostral 2 sections (p=0.014) and remained stable throughout the remainder of the rostral-caudal axis (Figures 2I, 2J).

#### D3R-only SPNs

We observed a distinct subpopulation of striatal D3R-only SPNs, albeit in far smaller numbers compared to D1R-only and D2R-only cells (Figures 2L-2O). The representative heat map analysis revealed small, dense pockets of D3R-only cells concentrated in the Shell and OT, primarily within the IC (Figure 2K). Unlike D1R- and D2R-only cells, there were no differences in the striatal distribution of D3R-only SPNs across either the dorsal-ventral or rostral-caudal axes (p>0.05; Figures 2L-2O, Figures S2I-S2L).

### Spatial distribution of striatal dopamine D_1_, D_2_, D_3_ receptor expression in co-expressing spiny projection neurons

Our multiplex RNAscope studies revealed several distinct striatal SPN subpopulations that co-expressed either DA D_1_ and D_2_ receptors (D1/2R), D_1_ and D_3_ receptors (D1/3R), or D_2_ and D_3_ receptors (D2/3R) (Figure 3, Figure S3). We found no evidence of SPNs that co-expressed all three DA receptors together, *i.e*., D1/2/3R. Threshold analyses ensured we did not double count cells (see Methods), enabling us to accurately distinguish SPNs that only expressed a single DA receptor from SPNs co-expressing multiple DA receptors. Quantification revealed that, while DA receptor co-expressing SPNs were present throughout striatum, these cells were found in lower numbers compared to SPNs that expressed single DA receptors. Interestingly, our analyses also demonstrated additional dorsal-ventral and rostral-caudal trends in cell distribution for the co-expressing neurons (Figure 3).

#### D1/2R SPNs

Quantification showed that D1/2R SPNs constituted ∼10-25% of the total pool of D1R^+^ and D2R^+^ cells in the striatum (Table S1). D1/2R SPN enrichment in the OT persisted throughout the rostral-caudal axis. Indeed, in rostral sections (Rostral 1-3), the OT possessed as much as double the D1/2R SPN density compared to all other striatal subregions (Figures 3B-3D) [Rostral 1: F(5,54) = 3.954, p=0.004; Rostral 2: F(5,65) = 4.227, p=0.002; Rostral 3: F(5,63) = 3.309, p=0.01)]. Moving caudally, however, we observed a gradual decrease in this OT enrichment (Figures 3B-3E). Closer inspection of the Shell subregions revealed similar trends across the rostral-caudal axis (Figure S3A-S3D).

Along the dorsal-ventral axis, we examined the relative proportion of D1/2R neurons out of the total number of SPNs that expressed either D1R or D2R. Our analyses confirmed the relative abundance of D1/2R SPNs in the OT compared to other striatal subregions. Indeed, OT D1/2R SPNs constituted 20-28% of the total number of D2R^+^ cells, while composing 12-16% of D1R^+^ cells in this subregion (Table S1). Besides the OT, we also observed changes in other striatal subregions along the dorsal-ventral axis. For example, in the Rostral 3 section, there was a significant increase in the proportion of D1/2R SPNs in the CPu versus the Shell relative to the total number of D1R^+^ cells (p=0.001). Additionally, in the most caudal section (Caudal), the proportion of Shell D1/2R SPNs relative to the total pool of D1R^+^ cells was significantly lower compared to other subregions including the CPu (p=0.014), Core (p=0.041), or OT (p=0.007).

We also found Shell-specific differences in relative densities of D1/2R SPNs along the rostral-caudal axis. Out of the total number of D1R^+^ cells, there were significantly more Shell D1/2R SPNs in the most rostral section (Rostral 1) compared to the most caudal section (Caudal) (p=0.011). In contrast, out of the total number of NAc shell D2R^+^ cells, there was a significantly higher proportion of D1/2R SPNs in the Rostral 2 section compared to Rostral 1 (p=0.046) (Table S1). Taken together, these data point to shifts in the distributions of co-expressing D1/2R SPNs across both dorsal-ventral and rostral-caudal axes, suggesting predefined patterns of receptor expression.

#### D1/3R SPNs

In contrast to the relatively small numbers of D3R-only SPNs, most D3R^+^ striatal SPNs co-expressed D3R alongside D1R (D1/3R SPNs). Nevertheless, compared to D1/2R SPNs, D1/3R SPNs constituted a much smaller proportion of the total D1R^+^ SPN population, ranging from 0-7% based on the striatal subregion and position along the rostral-caudal axis (Figures 3F-3J, Table S1). Along the dorsal-ventral axis, there was OT enrichment of D1/3R SPNs in the Rostral 1 section [F(3,58)=5.564, p=0.002; Figure 3G], with no further enrichment in more caudal sections (p>0.05; Figure 3H-J). Akin to D1/2R SPN distribution, the changes in D1/3R density along the rostral-caudal axis were exclusive to the Shell. Specifically, the proportion of Shell D1/3R SPNs out of total D1R^+^ neurons was higher in the Rostral 1 section compared to Rostral 2 (p=0.003), Rostral 3 (p=0.001), or Caudal (p=0.004) sections (Table S1).

#### D2/3R SPNs

Of the three DA receptor co-expressing SPNs, D2/3R SPNs represented the smallest striatal subpopulation (Figure 3K). Limited numbers of D2/3R cells were primarily expressed in the ventral striatum with no significant distribution in the CPu or Core. This pattern was present across all striatal sections throughout the rostral-caudal axis (Figures 3L-3O; Table S1).

Taken together, the OT represents with striatal subregion with the greatest enrichments in SPNs that co-express DA receptors. In fact, 11-15% of D1R^+^ cells co-expressed D2R and 20-26% of D2R^+^ cells co-expressed D1R in the OT (Table S1). While on a smaller scale, a large proportion of D3R^+^ SPNs co-expressed D1R in the OT, ranging from 3-7% (Table S1). Our spatial density heatmaps similarly showed that both D1/2R (Figure 3A) and D1/3R (Figure 3F) co-expressing SPN subpopulations were concentrated throughout the OT. Importantly, these maps also highlighted dense pockets of D1/2R and D1/3R co-expressing cells in the IC (Figures 3A, 3F). Consequently, we quantified the spatial distribution of these co-expressing subpopulations in the ventral striatum. We focused on the OT and IC to determine whether D1/2R and D1/3R cell subpopulations were preferentially enriched in one or both of these striatal subregions. We did not examine D2/3R neuron distribution since these cells were not significantly present in either ventral striatal subregion. We found that the D1/2R^+^ cells were preferentially enriched throughout the entire OT, whereas D1/3R co-expression was mainly restricted to the IC (Figure S4).

Overall, our multiplex RNAscope data led us to conclude that each striatal subpopulation exhibited distinct patterns of expression across dorsal-ventral and rostral-caudal axes (Figure 4). This 3D mapping also revealed gradients of differential distribution for several SPN subpopulations, particularly by DA receptor co-expressing cells.

**Figure 4.**
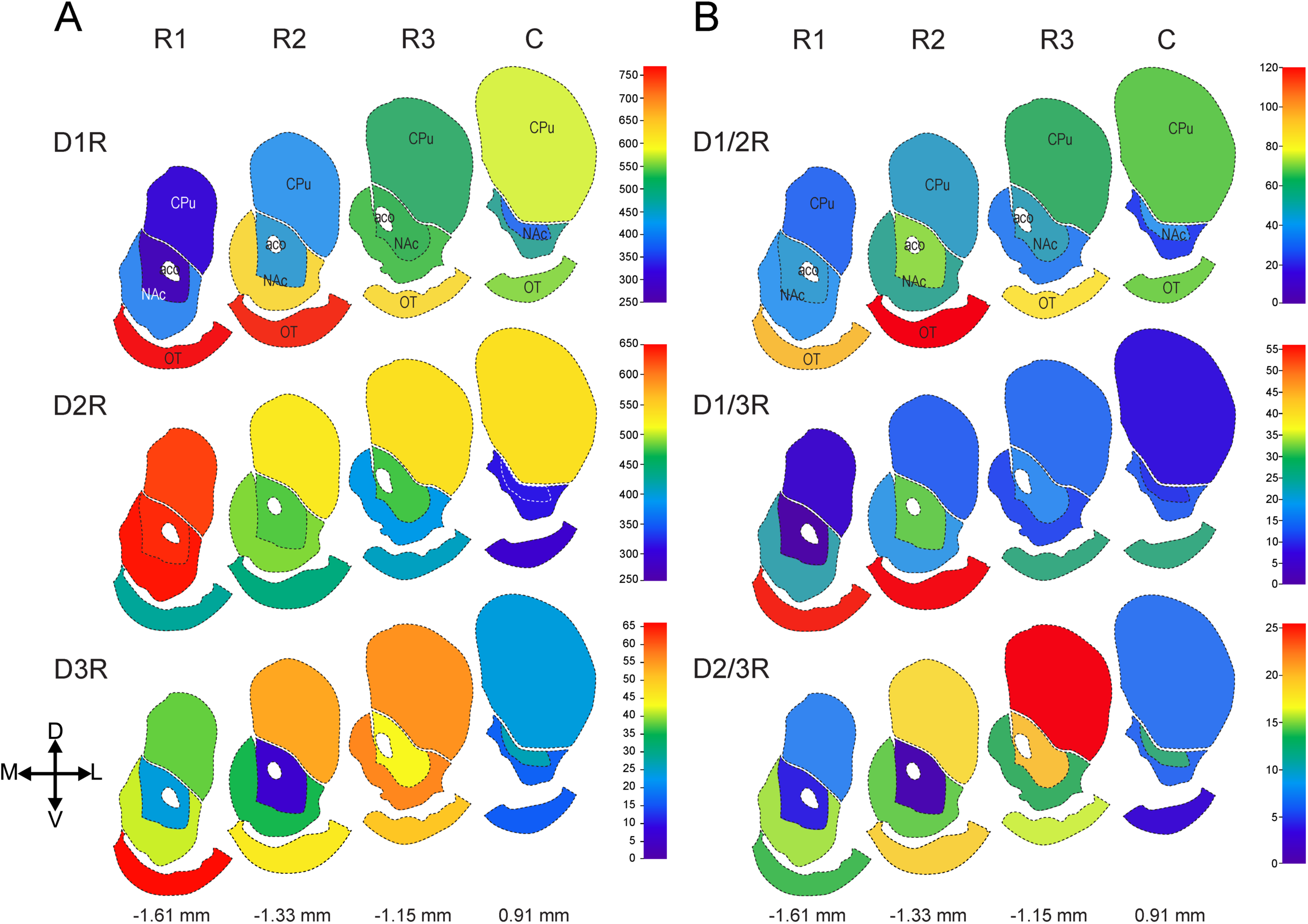
Schematic demonstrating 3D patterns of SPN DA receptor expression across dorsal-ventral and rostral-caudal axes. Distribution patterns of singly-expressing striatal SPNs expressing: D1R, D2R, or D3R **(A)** and SPNs co-expressing D1/2R, D1/3R, or D2/3R **(B)** in the caudate putamen (CPu), nucleus accumbens (NAc), anterior commissure (aco), and olfactory tubercle (OT). Changes in density were also assessed across a rostral-caudal axis (R1 = - 1.61mm, R2 = -1.33 mm, R3 = -1.15 mm, and C = 0.91 mm). The numbers on the scales represent the average positive cell density for each co-expressing receptor subpopulation. D=dorsal, V=ventral, L=lateral, M=medial.

### Sex differences in dopamine receptor expression within the Islands of Calleja

While there were no detected sex differences in most of the major striatal subregions, we discovered significant sex differences in the IC (Figure 5). Among the predominant striatal DA receptor subpopulations, the female IC was primarily composed of D1R-only cells compared to males, who exhibited significantly lower D1R-only cell densities (p=0.0011; Figure 5C). In contrast, the IC in male mice also contained D3R-only and D1/3R-co-expressing neurons, whereas female IC showed negligible density of these cells (D3R-only: p=0.0002, D1/3R: p=0.00089) (Figs. 5F, 5I). Interestingly, we found no evidence of D2R-only or D2/3R-only cells in the IC of either males or females (data not shown).

**Figure 5.**
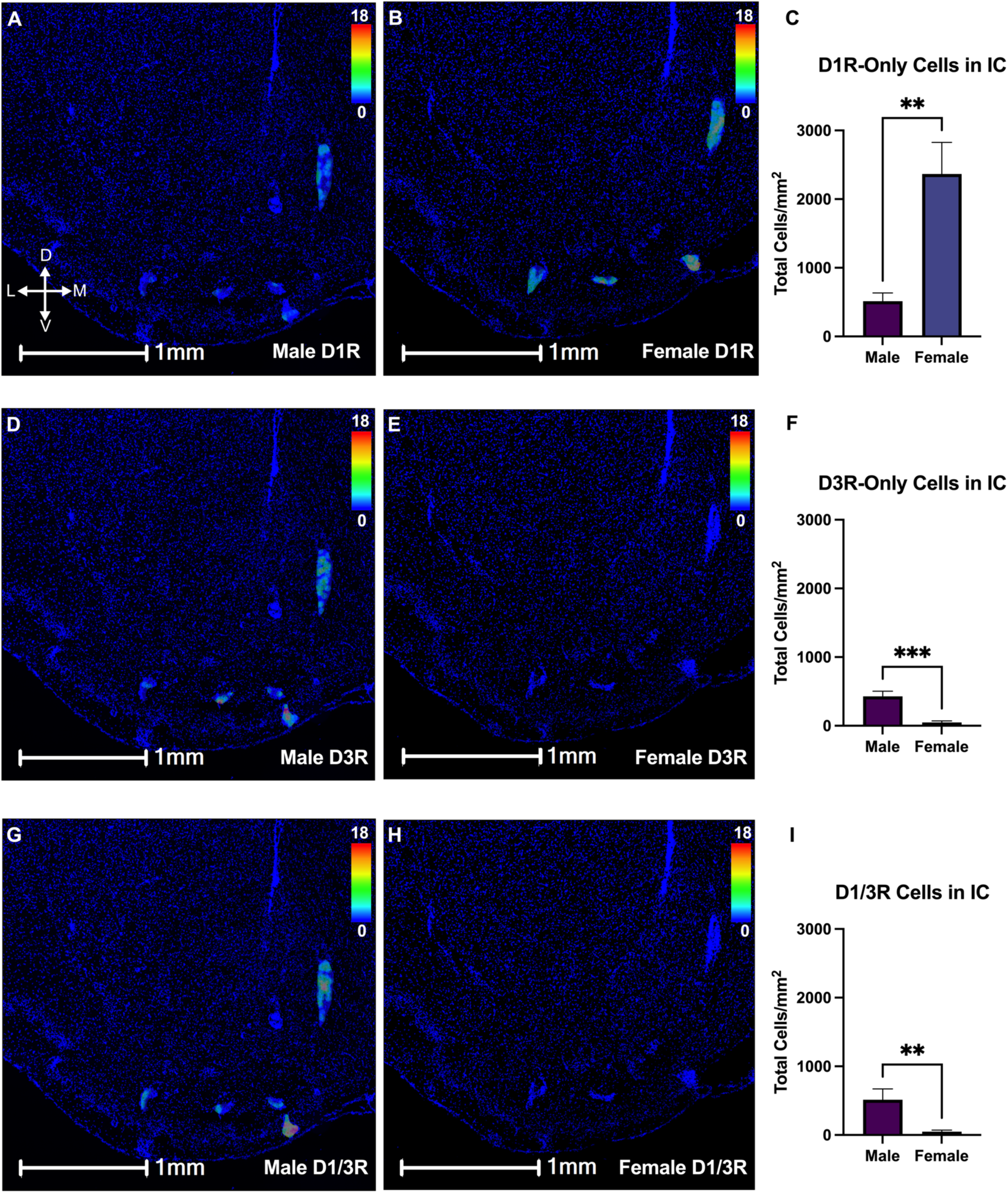
Sex differences in dopamine receptor expression within the Islands of Calleja (IC). **(A-C)** Representative images **(A, B)** and quantification **(C)** of D1R expression in the IC revealed that females primarily expressed D1R-only SPNS compared to males (**p=0.0011). **(D-I)** Representative images **(D,E,G,H)** and quantification **(F, I)** of D3R expression in the IC showed that the IC in males is also composed of D3R-only **(D-F)** and D1/3R+ **(G-I)** SPNs, unlike females who express negligible levels of either receptor subpopulation in this region (D3R-only: ***p=0.0002; D1/3R: **p=0.0089). For **Panels A, B, D, E, G, H**, scale bar = 1 mm; heat map legends represent the number of mRNA grains within a given cell. Mean ± SEM, n=3 male mice, n=3 female mice for all conditions. *p<0.05, **p<0.01, ***p<0.001. D=dorsal, V=ventral, L=lateral, M=medial.

### Conservation of distinct striatal cell type expression across multiple species

In parallel to our RNAscope mapping, we used single-nuclei multi-omics of the striatum across mammals to determine the molecular signatures unique to SPN subpopulations. While we showed that different subpopulations have distinct spatial distributions according to their respective DA receptor expression, RNAscope alone cannot simultaneously identify other genes that define these cells. On the other hand, the recent explosion of single-cell technologies including single-nuclei RNA sequencing (snRNA-seq) and single nucleus assay for transposase-accessible chromatin using sequencing (snATAC-seq), while incapable of providing spatial information, can determine whether striatal subpopulations are distinct according to their transcriptional and epigenomic profiles.

Despite their many advantages, it has been challenging to harmonize the reports of SPNs from single-cell studies in the last decade^46–48,51,53,54,58^. The studies varied by species, single-cell technologies, and sampled striatal subregions. Furthermore, each study used different naming conventions for striatal cell types, making it difficult to disentangle the SPN heterogeneity beyond the traditional D_1_-like versus D_2_-like dichotomy (Figures 6A, 6B). To address this, we conducted a comprehensive comparative analysis using eight publicly available single-cell datasets to consistently define SPN diversity. We selected datasets that spanned: 1) multiple mammalian species (mouse, rat, rhesus macaque, and humans), 2) snRNA-seq or snATAC-seq, and 3) the original sources of distinct annotation use by other studies.

**Figure 6.**
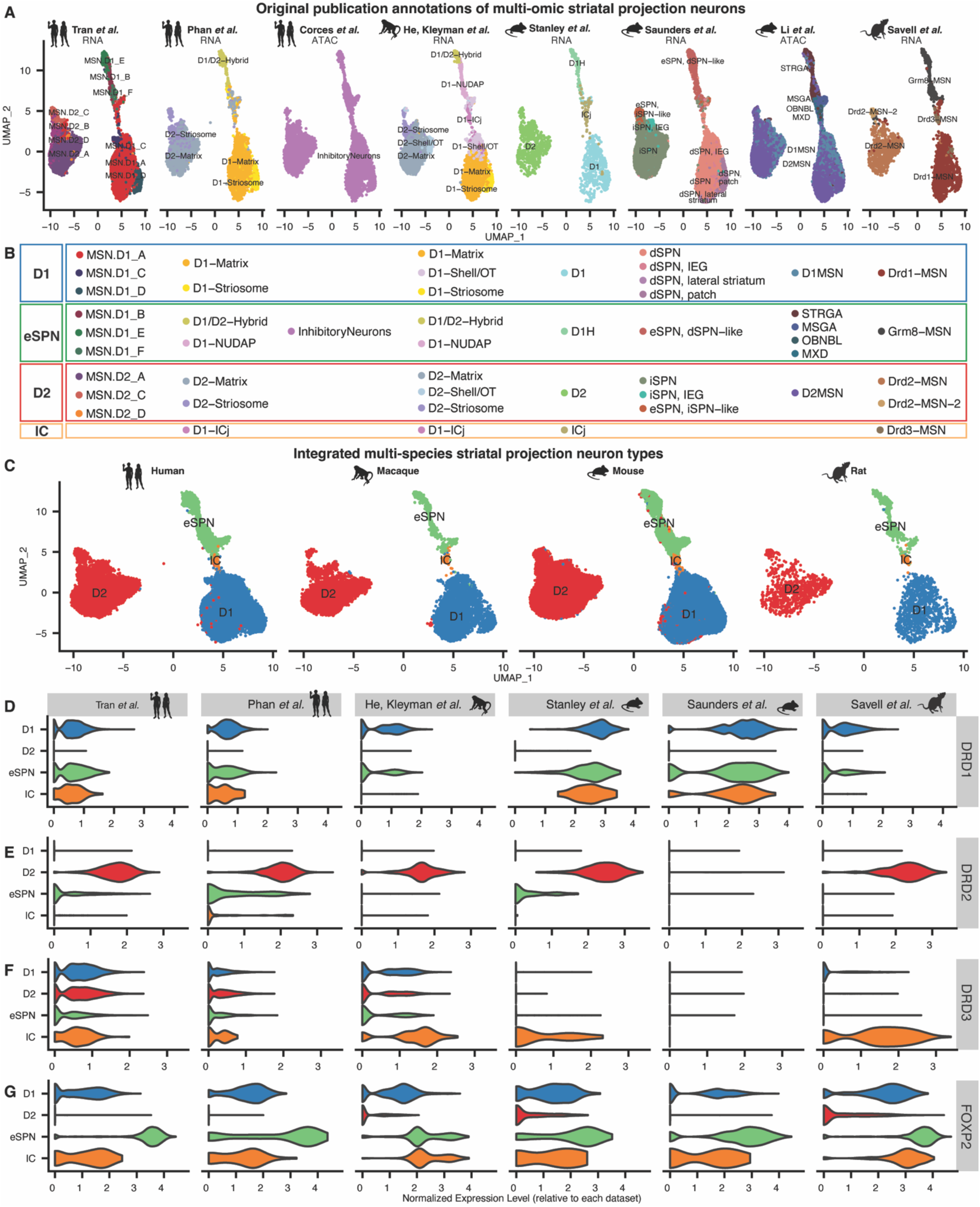
Multi-species consensus of distinct striatal projection neuron cell types. **(A, C)** UMAP projections of multi-species striatum datasets using the original publication labels to identify cell type clusters **(A). (B)** Consolidation of SPN annotations from the original publications into a parsimonious consensus. **(C)** UMAP projections of the integrated, multi-species cell type consensus labels. **(D-G)** Violin plots of normalized relative expression for indicated genes within each dataset stratified by consensus labels.

We established an integrated dataset to label SPNs parsimoniously across multiple species and datasets. Despite methodologic differences between datasets (Figures S5A, S5B), we successfully clustered SPNs between all datasets without dataset- or sample-specific biases (Figure S5C). When integrated together, we found that the same four SPN cell types were conserved across rodents and primates: D1R-only, D2R-only, eccentric SPNs, and IC neurons (Figure 6C). Although some cell types were named differently by their respective studies (Figure 6B), we reconciled their respective annotations. Known dimensions of SPN spatial organization, such as striosome versus matrix, were also inconsistently annotated. We therefore merged the multi-omics data with the spatial distributions of these cell types based on our previous RNAscope data^46–48,51,52^ to add spatial “subtypes” (*e.g.*, D1-striosome, D1-matrix, etc.) (Figure S5D).

We identified two distinct SPN cell types that co-expressed DA receptors beyond the canonical dichotomy: one cell type unique to the IC and another cell type previously annotated as eccentric SPNs or ‘eSPNs’^46^. These cell types mapped onto our D1/3R and D1/2R co-expressing cells, respectively. eSPNs possessed unique transcriptomic profiles and spatial distributions, distinguishing them from D1R-only, D2R-only, or IC neurons (Figure 6C). We confirmed the normalized expression patterns of DA receptors across neuronal subtype clusters. *DRD1* was highly expressed in D1R-only, eSPN, and IC neurons (Figure 6D). In contrast, *DRD2* expression was strongest in D2R-only neurons with measurable, but weaker expression in eSPNs (Figure 6E). *DRD3* expression inconsistently marked IC neurons across datasets. IC neurons possessed the highest expression of *DRD3* in the mouse, rat, and rhesus macaque datasets^47^ (Figure 6F), yet human IC neurons did not express *DRD3* more than other cell types.

Next, we assessed whether eSPNs and IC neurons co-expressed multiple DA receptors in snRNA-seq datasets. We confirmed DA receptor co-expression with a scatter plot of *DRD1, DRD2,* and *DRD3* in single cells (Figure S6). We found eSPNs: 1) co-expressed *DRD1* and *DRD2*, and 2) their DA receptor co-expression was conserved across species (Figure S6A). IC neurons also co-expressed *DRD1* and *DRD3*, but the co-expression was less distinct compared to D1R-only SPNs (Figure S6B). From the cross-species single-cell RNA-seq data, we were only able to quantify the proportion of SPNs across gross dorsal-ventral gradients (Figure S5E). We found stable trends in the relative proportions of D1R-only and D2R-only SPNs that were consistent across species. In the dorsal striatum, D1R-only and D2R-only SPNs constituted ∼90% of total SPNs with 45% of each type represented by D1R-only and D2R-only SPNs, respectively; eSPNs and IC neurons made up the remaining 10% of SPNs. The proportions of eSPNs and IC neurons in ventral striatum were greater than in the dorsal striatum (Figures 3A and 3B, respectively). This was consistent with our RNAscope mapping which provided finer spatial resolution of the same SPNs. These cross-species transcriptomics data therefore both strongly support and complement our RNAscope mapping results.

A common finding across the single-cell multi-omics data was that the various striatal cell types were distinctly marked by other genes beyond DA receptors. Therefore, we computed a list of differentially expressed genes between cell types that were conserved across species and datasets. This allowed us to establish a resource for species-conserved markers of unique striatal cell types (Table S2). We discovered that D1R-only neurons were marked by 43 genes (FDR < 0.05) including *DRD1*, *PDYN*, and *TAC1* (FDR < 1.6e-79), well-established markers for these neurons. D2R-only neurons were marked by 55 genes (FDR < 0.05) including *DRD2* and *PENK* (FDR < 2.0e-188). In contrast, eSPNs were marked by a different set of 54 marker genes, none overlapping with D1R-only or D2R-only markers, establishing eSPNs as a transcriptomically distinct cell type^46–48,52,58^. Top marker genes for eSPNs included *FOXP2* and the previously reported eSPN marker, *CACNG5* (FDR < 1.7e-29)^46^. In addition, we identified numerous marker genes of SPNs at the spatial subtype level, further providing a molecular basis for the spatial organization of distinct SPN subtypes (Table S2). Finally, IC neurons were marked by only 4 conserved marker genes across species, with only 1 overlapping D1R-only neurons (FDR < 0.05) and none in caudoputamen. However, this could be due to the lower representation of IC neurons in NAc/OT datasets and limited representation in caudate/putamen datasets (Figure 6C).

### Spatially distinct localizations of eSPNs throughout the dorsal and ventral striatum

Inconsistent annotation of eSPNs from snRNA-seq data has resulted in conflicting descriptions of this cell type (Fig 6A, 6B). Therefore, we used the eSPN marker, *FOXP2*, to link snRNA-seq results to our RNAscope mapping. According to our snRNA-seq data, *FOXP2* was expressed in eSPNs at significantly higher levels compared to other striatal cell types in a species-conserved manner (log_2_ fold-change = 2.5, FDR = 1.8e-34; Figure 6G). We experimentally validated this enrichment with multiplex RNAscope, labeling *Drd1*, *Drd2*, and *Foxp2* in the adult mouse striatum. We confirmed that *Foxp2* labeled the D1/2R co-expressing neurons across the striatum (Figures 7A-7G). In the dorsal striatum, *Foxp2* was expressed widely in D1R-only neurons and eSPNs with substantially less *Foxp2* expressed in D2R-only neurons (Figure 7G). Quantitation of these RNAscope data revealed that dorsal striatal eSPNs expressed a moderately higher level of *Foxp2* mRNA grains per cell compared to D1R-only cells (eSPN: 5.18 ± 0.39 grains per cell, D1R-only: 3.95 ± 0.042: Cohen’s D = 0.50). eSPNs expressed largely more *Foxp2* versus D2R-only SPNs (D2R-only: 2.54 ± 0.17 grains per cell: Cohen’s D = 1.0). We also established that eSPNs were evenly distributed across the dorsal striatum according to *Foxp2* expression (Figure 7E). This is consistent with our above data (Figure 3A) and previous studies that relied on other eSPN makers like *Otof*, *Cacng5*, and *Casz1*^46^. In the ventral striatum, there was only a small difference of *Foxp2* expression in eSPNs versus D1R-only SPNs (eSPN: 6.74 ± 0.53 grains per cell, D1R-only: 7.11 ± 0.088: Cohen’s D = -0.49). Ventral striatal eSPNs still expressed substantially more *Foxp2* compared to D2R-only SPNs (D2R-only: 1.52 ± 0.17: Cohen’s D = 1.05). Furthermore, *Foxp2*^+^ cells formed well-defined clusters throughout the OT and NAc (Figures 7E, 7F), consistent with the D1/2R clusters featured in Figure 3A and in previous studies^47^. We additionally confirmed our *Foxp2* results with RNAscope studies of *Casz*1 expression, finding a similar dorsal and ventral striatal distribution (data not shown). Importantly, our experimental RNAscope findings quantitatively recapitulated the relative cell type-specific differences in *FOXP2* expression from multi-species snRNA-seq data. These expression differences allowed us to further define eSPNs, D1R-only, and D2R-only neurons according to their respective levels of *FOXP2* expression (Fig 6F, 7C).

**Figure 7.**
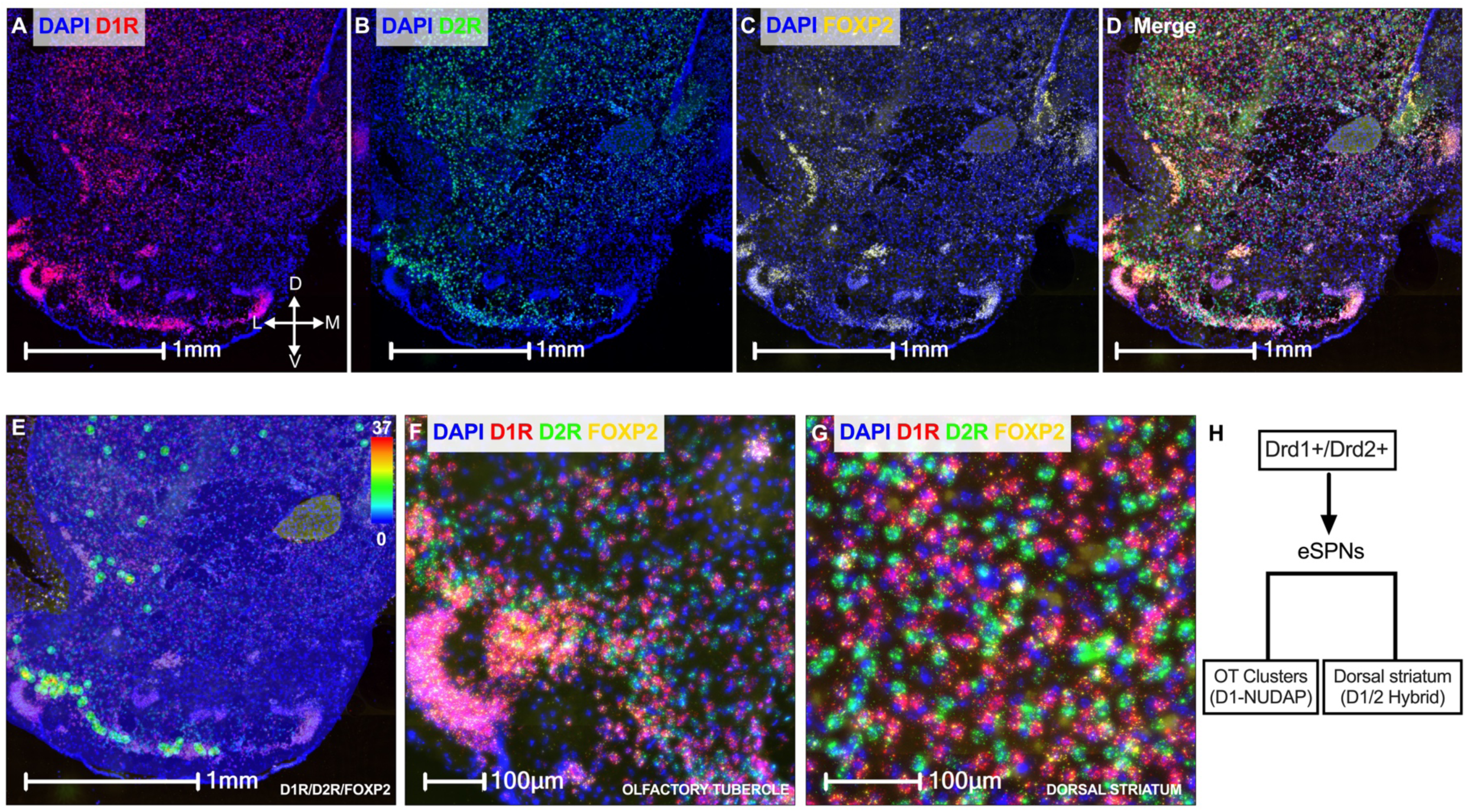
mRNA expression of Foxp2 in dorsal and ventral striatum. **(A-D)** Representative multiplex RNAscope images of D1R (A), D2R (B), Foxp2 (C), and merged channels (D). **(E)** Spatial density heatmap of cells positive for the co-expression of D1R, D2R, and Foxp2. **(F)** Magnified image of the OT, highlighting dense clusters of D1R/D2R/Foxp2 co-expression. **(G).** Magnified image of the CPu, displaying distributed co-expression of D1R/D2R/Foxp2. **(H)** Schematic defining D1/D2R^+^ neurons that co-express high levels of Foxp2 as eSPNs. These eSPNs are further subcategorized on the basis of their striatal localization, either clustered in the OT (NUDAP) or distributed throughout the striatum. D=dorsal, V=ventral, L=lateral, M=medial.

Our multi-omics analyses revealed different patterns of eSPN localization. Data from He, Kleyman *et al.* suggested that eSPNs divided into two subtypes, D1/2 Hybrid and D1-NUDAP (D1-Neurochemically Unique Domains in the Accumbens and Putamen) according to their spatial distributions^48^ (Figures 6A, S5D). Consistent with this, our RNAscope data (Figures 3A, 7E) showed that D1/2R co-expressing neurons (labeled as eSPNs), had two patterns of localization between the dorsal and ventral striatum in mice. The first spatial pattern featured evenly distributed D1/2R SPNs amongst D1R-only and D2R-only SPNs in the dorsal striatum (Figure 3A, 7G). This corresponded to the spatial distribution of D1/D2 Hybrid neurons in the rhesus macaque caudate and putamen. In contrast, the second pattern of D1/2R SPNs showed distinct cell clusters within the OT and NAc (Figure 3A, 7F). This also corresponded to the D1-NUDAP neurons reported in the rhesus macaque OT. Overall, this convergence of spatial and transcriptomic data across multiple species and striatal subregions leads us to conclude that eSPNs comprise two spatially distinct subtypes (Figure 7H).

### Striatal cell types in human disease risk

To better understand the roles of specific striatal cell types in human disease, we examined whether the epigenomic profiles of SPN cell types intersected with human genetic markers of neurological and psychiatric disease. Previous studies showed enrichment of striatal neurons with human disease, albeit at lower cell type resolution or with indirect approaches via non-human data or with large neighborhoods surrounding marker genes that gloss over the locations of genetic risk variants^54,59–61^. Instead, we used our SPN labels of snATAC-seq of postmortem human caudate^54^ to specify the epigenomic regions of which cell types drive the genetic associations with human disease traits. We performed a conditional hereditability enrichment analysis to separate the specific contributions of each striatal cell type. We intersected these profiles with a panel of 64 human traits spanning multiple disease and psychosocial domains (*e.g.*, neurodegeneration, psychiatric disorders, substance use, psychosocial domains, sleep, and non-brain related negative control traits). We discovered that D1R-only SPNs were enriched in cocaine-dependent risk variants^62^ independently of other striatal cell types (*e.g.*, D2R-only, eSPN, interneurons, glia, etc.; Figure 8). D2R-only SPNs were enriched in 3 psychiatric traits (major depressive disorder, schizophrenia, cross-psychiatric trait)^63–65^ and educational attainment^66^. eSPNs, in contrast to D1R-only and D2R-only SPNs, significantly contributed to more than twice the number of human disease traits and pyschosocial domains. This included psychiatric (cross-psychiatric trait, bipolar disorder, schizophrenia)^63,65,67^, substance use (current versus non-smoker)^68^, numerous psycho-social domains (BMI, education, risk tolerance)^66,69,70^, and sleep (morning versus evening preference)^71,72^. Interneurons and glial cell types were also enriched in several neuropsychiatric traits. The complete numerical results are reported in Table S3.

**Figure 8.**
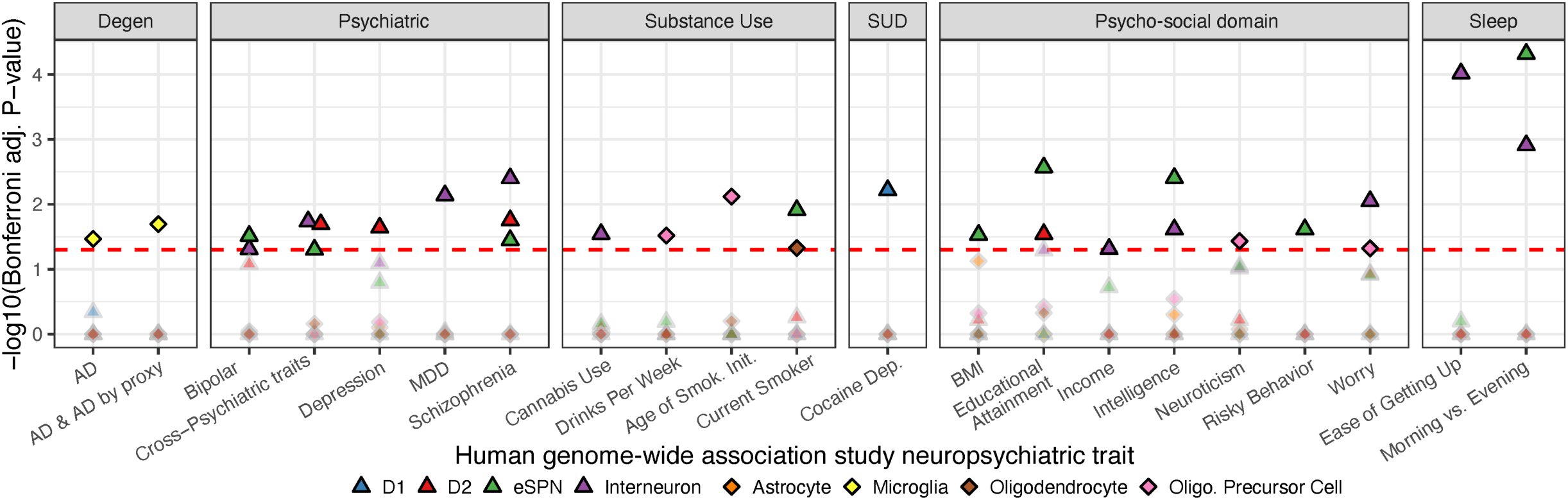
Unique contributions of striatal cell types to genetic risk of neuropsychiatric diseases. Scatter plot of significance from conditional hereditability enrichment analyses of striatal neuron and glial ATAC-seq profiles with a GWAS panel of human neuropsychiatric traits. GWAS traits are grouped by general disease processes or psychosocial domains. Each point represents a tested cell type-trait combination, and the shape and color represent different neuronal or glial cell type. The complete numerical results are reported in Table S3. The dotted line marks the P_Bonf_< 0.05 cut off. Significant associations are plotted as bold, black outlined points. Non-significant associations are plotted as gray outlined points with transparency. AD: Alzheimer’s Disease, BMI: Body-mass index, Degen: neurodegenerative disease, Dep: dependence, Init: initiation, MDD: Major Depressive Disorder, SUD: substance use disorder.

## Discussion

There are well-defined relationships between striatal anatomy and its variety of functional roles^73–76^. For example, functional differences have been described for distinct NAc subregions where DA transient amplitude and frequency differ in Core versus the Shell^77^. Furthermore, in the dorsal striatum, neuronal responses to different sensory modalities follow a medial-lateral gradient^78^. Regional differences in DA signaling have also been extensively documented in the context of drugs of abuse where the dorsal versus ventral striatum exhibits different molecular responses in drug-seeking behavior^79,80^. What signaling mechanisms can explain these differences? Traditionally, striatal DA signaling has been divided along a functional dichotomy between D1R^+^ and D2R^+^ SPNs. In fact, at a basal level, D2R^+^ SPNs exhibit increased excitability and higher spontaneous excitatory postsynaptic currents compared to D1R^+^ SPNs^15^. In the context of drugs of abuse like cocaine and opioids, responses in D1R^+^ and D2R^+^ SPNs are heterogeneous, raising the possibility of unique subpopulations of striatal cells^22,81^. Indeed, SPNs that express multiple DA receptor subtypes are increasingly implicated in a variety of contexts, including reward-learning, decision-making, and goal-directed movement^22,82,83^.

Our use of multiplex RNAscope provides direct confirmation of multiple SPN subpopulations in spatially well-defined patterns. Using this approach, we established a definitive, comprehensive 3D map of DA receptor-expressing SPN subtypes in adult mouse striatum. We found unique subpopulations of SPNs that co-express either D1/2R, D1/3R, or D2/3R alongside SPNs expressing D1R, D2R, or D3R alone. These different SPN subpopulations were heterogeneously distributed throughout the striatum along dorsal-ventral and rostral-caudal axes. This is consistent with prior spatial transcriptomic studies employing that similarly demonstrated several transcriptionally diverse SPN subpopulations that were organized into distinct spatial domains within the striatum^84,85^. While these approaches have provided critical new insights into gene expression patterns across space, they have important limitations including circumscribed numbers of genes that can be studied over time and resolutions that current advances in neurophysiology have yet to effectively investigate. Furthermore, though previous work deeply profiled the expression patterns of SPNs, these studies annotated SPN subpopulations independently, resulting in a lack of consensus as to how these SPN cell types should be defined. To rectify this, we conducted a multi-species, multi-omic integration of eight independent single-cell datasets. Our synthesis supports a simple classification of four SPN subpopulations: D1R-only, D2R-only, eSPN, and IC. These subpopulations were conserved from rodents to primates, and reliably defined across anatomic, transcriptomic, and epigenomic domains. Furthermore, our integration of RNAscope mapping with multi-omic analyses connected the D1/2R and D1/3R co-expressing SPNs to the eSPN and IC cell types, respectively. The epigenomic profiles of the different SPN cell types contributed uniquely to genetic risk for complex human disease traits. Strikingly, we found that minority cell types like eSPNs were disproportionately enriched in numerous disease and psychosocial traits compared to more populous striatal cell types. Collectively, we have established an integrated profile of distinct SPNs defined by anatomy, transcriptomics, and epigenomics. Moreover, this SPN heterogeneity offers a new understanding of striatal neuron organization, function, and relationship to human disease.

Consistent with evidence of SPN heterogeneity, we have demonstrated the existence of several SPN subpopulations that co-express either D1/2R, D1/3R, or D2/3R in addition to the established populations of D1R-only and D2R-only SPNs. Though these DA receptor co-expressing SPN subpopulations constitute a minority of the total striatal SPN pool, we consistently observed DA receptor co-expression in specific striatal subregions in well-defined 3D patterns. Our data also corroborate and expand upon earlier work which similarly identified discrete SPN subpopulations along a dorsolateral-ventromedial gradient^47^. We also found that D1/2R SPNs were enriched in the OT compared to other striatal regions. Though the precise functions of dopaminergic signaling OT are still unclear, earlier studies demonstrated that D1R and D2R play opposing roles in functions mediated by OT, including in odor-conditioned reward responses^86,87^. D3R functions in the IC are similarly poorly understood. However, clues come from findings that D3R^+^ SPNs are enriched in the IC and play a critical role in grooming behavior in mice^88^.

The co-expression of different DA receptors in the same SPNs raises a key question: does DA receptor co-expression confer unique properties compared to expression of either receptor alone? Computational models proposed that D1/2R-co-expressing SPNs have distinct roles in reward-punishment-risk-based decision-making^82^. In the disease context, D1/2R SPNs are uniquely affected by dopaminergic deafferentation in preclinical models of Parkinson’s disease^89^. However, the mechanisms by which D1/2R co-expression relates to function remain poorly understood. Prior work suggested that co-expression of D1R and D2R leads to the formation of higher-order heteromeric complexes^90^. These D1/2R complexes may enable recruitment of downstream effector molecules (*i.e.*, Gα_q_) that are different from those recruited to either D1R (Gα_s_) or D2R (Gα_i/o_) alone^91^. Functionally, D1/2R receptor heteromeric complexes have been implicated in cocaine-induced locomotion and self-administration^92^; these heteromers are also associated with anxiety-like and depressive-like behaviors^93^. Outside of the brain, D1/2R heteromers may also modulate pancreatic islet function^94^. However, the physiological relevance of DA receptors heteromers remains controversial as most of these studies relied on exogenous overexpression. This raises the possibility that D1/2R heteromers may not be relevant physiologically since endogenous levels are often far lower than the levels produced via heterologous overexpression. We therefore posit that, even in the absence of endogenous D1/2R heteromers, co-expression of D1R and D2R is physiologically relevant. Concurrent activation of D1R and D2R in D1/2R SPNs may result in downstream signaling changes that are intermediate between stimulatory D1R^+^ and inhibitory D2R^+^ SPNs.

We also identified D1/3R co-expression primarily in the NAc ventromedial shell and IC. It has been suggested that D1/3R co-expression serves as a buffering mechanism to temper the stimulatory signaling by D1R and inhibitory signaling by D3R^95^. Functionally, D1/3R signaling is relevant to levodopa-induced behavioral sensitization in hemi-parkinsonian rodents and has been hypothesized to play a role in the actions of antipsychotic medications^95^. Studies to dissect the respective roles of D1/2R and D1/3R subpopulations have been limited due to the challenges in selectively targeting these subpopulations. Of the studies that have attempted to examine these co-expressing subpopulations, many have either been *in vitro*, relied on ectopic receptor overexpression, or employed bacterial artificial chromosome (BAC) transgenic mice expressing fluorescent tags in D1R^+^ versus D2R^+^ SPNs, leading to conflicting findings^96–98^.

The combination of our 3D mapping of D1R, D2R, and D3R expression across dorsal-ventral and rostral-caudal axes and multi-omics analyses of the striatum revealed gradients of differential distribution for several SPN subpopulations, especially for co-expressing cells. This is best exemplified in D1/2R SPNs which exhibited a gradient of cell density that increased along the dorsal-ventral axis while decreasing along the rostral-caudal axis. What is the relevance of such gradients? Gradients are well-established features of brain development, including the striatum^99^. Indeed, gradients of signaling by morphogens such as sonic hedgehog (Shh) and Wnts are critical for striatal development and dorsal-ventral patterning^99,100^. These gradients may also persist throughout adulthood^101^. We posit that spatial patterns of co-expressing SPNs represent echoes of earlier developmental programs, reflecting persistent morphogenic gradients that gave rise to these specific subpopulations. Numerous SPN marker genes are neurodevelopmentally-regulated transcription factors that coordinate gene expression across fetal development, neuronal migration, differentiation, and maturation^102^. Moreover, different interneuron subpopulations are similarly organized according to gradients in both mouse and human striatum^103^. For example, *PTHLH*-expressing parvalbumin^+^ interneurons, which possess distinct electrophysiological properties, are distributed along a gradient in the dorsal striatum^103–105^. This suggests that different SPN subpopulations may themselves be subject to region- and state-dependent regulation by distinct combinations of interneurons along the striatal gradients^103,106^.

The cell types that we have described are well conserved across species and are reproducibly detected in multiple independent reports^45–53^. However, due to the ongoing evolution of multi-omics technologies, the sequencing depth of the datasets we analyzed is variable. For example, the number of unique transcripts per cell and genes per cell from the Saunders *et al.* mouse dataset, which preceded other datasets, was insufficient to directly measure *Drd2* and *Drd3* in the D2 and IC populations. Nevertheless, Saunders and colleagues were still able to report a D2R-only population that clustered apart from other SPN cell types using other transcripts^46^. Though we detected low but consistent levels of D1/2R and D1/3R co-expressing cell types in the single-cell data, we did not find a distinct D2/3R co-expressing SPN population. We believe that limits of single nuclei RNA-seq technologies preclude us from reliably measuring D2/3R co-expressing neurons, in contrast to our detection of this cell type by RNAscope at the single-cell level. Yet, despite these differences, we find one-to-one correspondence of our RNAscope mapping with the integrated single-cell data. The convergence between spatial mapping and multi-omics also supports a simplified classification of SPN cell types defined by their expression of DA receptors (D1R-only, D2R-only, D1/2R, D1/3R) and anatomic localization. Indeed, canonical D1R-only and D2R-only expressing cell types are known to have striosome and matrix subtypes^107–111^. Our combined multi-dimensional data defined the D1-NUDAP subtypes as D1/2R co-expressing eSPNs clustered within the OT^48^. Similarly, D1/D2 Hybrid subtypes are D1/2R co-expressing eSPNs evenly distributed throughout the striatum^48^. The D1-NUDAP subtype was previously defined in human and non-human primate brain using radiolabeled DAMGO binding assays to target the mu-opioid receptor^112,113^. We await future work to ascertain whether the two eSPN subtypes indicate functionally distinct circuits.

Finally, our conditional hereditability enrichment analysis identified associations between several striatal cell types to a broad panel of human neurological and psychiatric diseases as well as several psychosocial domains. Our heritability enrichment analyses are consistent with previous studies that linked SPNs to human diseases^54,59–61^. Notably, these earlier studies were limited by the resolution of their annotations since they did not differentiate SPN cell types. On the other hand, the ability to now separate SPNs into discrete cell types allowed us to resolve their contributions to human disease traits, providing new therapeutic targets. Using our analyses, we found significant enrichment of D1R-only SPNs with cocaine dependence risk variants. This is consistent with work linking D1R-only SPNs to cocaine exposure and reward^22,52,114–119^. Similarly, we recently connected D1R signaling to incubation of cocaine craving where mice exhibited decreased membrane excitability of D1R-expressing, but not D2R-expressing SPNs within the NAc shell after withdrawal from cocaine self-administration^120^. Likewise, different subtypes of D1R SPNs modulate cocaine reward and addiction in mouse models^121^. D2R-only SPNs were significantly associated with psychiatric traits including major depressive disorder and schizophrenia which is in line with an extensive literature connecting D2R signaling with both illnesses^4,12,122^. Indeed, antipsychotic medications, which block striatal D2R, are first-line treatments for schizophrenia and are increasingly used to treat mood disorders as well^123–125^. Importantly, compared to D1R-only and D2R-only SPNs, eSPNs were disproportionally overrepresented in these analyses, suggesting that this cell type is functionally distinct and plays critical roles in healthy and disease conditions. Further work is clearly needed to elucidate the biological mechanisms underlying these associations between the different disease states and the respective striatal cell types.

Overall, our study provides a definitive characterization of DA receptor expression in adult striatum. This reveals a much more heterogenous population of SPNs than previously appreciated and includes several distinct subpopulations of co-expressing striatal neurons. Our findings open the door to better understanding the functional relevance of these unique cell populations in the contexts of healthy and disease states.

## Methods

### Animals

Adult C57BL/6J mice from The Jackson Laboratory (stock 000664, 8-14-wks-old; Bar Harbor, ME) were used for this study. All animals were group-housed in a 12/12 light cycle with water and rodent chow provided *ad libitum*. All procedures were performed in compliance with ARRIVE guidelines^126,127^ and the Institutional Animal Care and Use Committee at the University of Pittsburgh (protocol #22071493).

### Brain tissue preparation

Mice were humanely euthanized followed by rapid brain collection and snap-freezing in isopentane; brains were then stored at -80°C until use. Brains were sectioned using a cryostat at -18°C. 14μm sections were collected at bregma -1.61mm (termed Rostral 1), bregma -1.33mm (termed Rostral 2), bregma -1.15mm (termed Rostral 3), bregma +0.91mm (termed Caudal) (Figure 1). Following mounting on slides (Superfrost+, Fisher Scientific, Waltham, MA), samples were first dried at -20°C for 2h, followed by additional drying at -80°C overnight.

### Multiplex fluorescent *in situ* hybridization

Multiplex RNAscope technology (ACD Bio, Neward, CA) was used to profile mRNA expression of *Drd1a*, *Drd2*, and *Drd3* as well as *Foxp2* and *Casz1* genes. RNAscope labelling of fresh frozen brain tissue was conducted according to manufacturer instructions using the ACD Bio RNAscope v1 kit. Briefly, pre-mounted tissue sections were post-fixed on slides in pre-chilled 4% paraformaldehyde (4°C, 60 min). Tissue underwent dehydration in successive ethanol baths of increasing concentration (50%, 70%, 100%). After drying, a hydrophobic barrier was drawn around samples, followed by incubation with probes (2h, 40°C) to detect cell-specific mRNA expression of mouse *Drd1a* (ACD Bio, Mm-Drd1a, Cat. 406491), *Drd2* (Mm-Drd2-C2, Cat. 406501-C2), *Drd3* (Mm-Drd3-C3, Cat. 447721-C3), *Foxp2* (Mm-Foxp2, Cat. 428791), and *Casz1* (Mm-Casz1-C3, Cat. 502461-C3). Following hybridization, slides underwent signal amplification with the respective probes allocated to each channel labeled with the following fluorophores: Channel 1, Atto 550; Channel 2, Alexa fluor 647; Channel 3, Alexa fluor 488. Tissue was then counter-stained with DAPI to label cell nuclei and mounted with Vectashield (Vector Labs, Neward, CA). Slides were stored at 4°C until imaging.

To test signal specificity of the RNAscope probes in our striatal tissue samples, we used RNAscope 3-plex Positive Control Probe (ACD Bio, Cat. 320881) and 3-plex Negative Control Probe (ACD Bio, Cat. 320871). Tissue preparation, hybridization, and signal amplification were as described above. The species-specific 3-plex Positive Control Probe labeled housekeeping gene controls for all 3 channels: Channel 1: DNA-directed RNA polymerase ll subunit RPB1 (POL2RA); Channel 2: Cyclophilin B (PPIB); and Channel 3: Ubiquitin C (UBC)(Figure S1A, B). The universal 3-plex Negative Control Probe contained probes targeting the DapB gene for all 3 channels (Figure S1C, D).

### Fluorescence slide scanning microscopy

Fluorescence imaging of labeled slides was performed using the Olympus VS200 automated slide scanner (Olympus, Center Valley, PA). Exposure times were adjusted for each fluorophore to ensure proper signal distribution with no signal saturation. Coronal sections were identified by experimenters and Z stacks of 3μm (3 Z-planes spaced by 1μm each) were imaged using a dry 20X objective lens (N.A. 0.8). Identical exposure settings and magnifications were consistently applied to all slides. After image acquisition, Z stacks were converted to two-dimensional maximum intensity projections and deconvolved using NIS Elements software (Nikon, Melleville, NY).

### Image analysis

Image analysis was performed using the HALO image analysis platform equipped with a fluorescent *in situ* hybridization module (Version 3.0, Indica Labs, Albuquerque, NM). Nuclei were quantified as DAPI-stained objects with the minimum cytoplasmic radius set at 5μm. Puncta corresponding to the respective mRNA probes were quantified as any 0.03-0.15μm^2^ object. We established a methodology to define cells specifically labelled by the respective probes above non-specific background. 1) We normalized the number of puncta per probe by dividing by the number of mRNA grains by total area analyzed (grains/area analyzed). 2) We then tested thresholds of 3x, 5x, 7x, and 10x above the puncta/area analyzed values based on our earlier RNAscope studies^128^. Examining the grains/area analyzed at 7-fold above the baseline (7x threshold) best optimized signal-to-noise in our datasets, providing accurate identification of appropriately labelled cells while minimizing non-specific signal. 3) We quantified the number of cells that reached this 7x threshold for each respective probe in a given sample. 4) To calculate the cellular density of positive cells, we divided the number of positive cells by the total area analyzed and converted to mm^2^ units.

To generate heat maps showing distribution of cells positive for single or double expression of dopamine receptors, images were analyzed using the spatial analysis module on the HALO platform. The previously described thresholds were also applied to account for background in the heat maps.

### Multi-species single-nucleus comparative transcriptomics and epigenomics

To harmonize annotations of striatal projection neuron subtypes across species and datasets, we analyzed eight published, annotated datasets in an unbiased manner, representing the broadest overview of single-nuclei RNA-seq and ATAC-seq data sampled from each species: mouse^45–47,53^, rat^51^, rhesus macaque^48^, and human^49,50,54^. The inclusion criteria for these comparative analyses are: 1) publicly availability of the data, 2) the data are annotated, 3) inclusion of single-cell or nuclei RNA-seq or ATAC-seq approaches, 4) from any striatal subregion, and 5) from a mammal. From each study, we focused the cell type integration at the level of projection neurons rather than glia or sparse interneurons inconsistently sampled across datasets. For the Phan *et al.* and Savell *et al.* datasets, we only included subjects in the unaffected comparison or saline injection group, respectively.

To map orthologous cell types at single-cell resolution across species and datasets, we performed integration analyses that were anchored using human gene identifiers. We subset genes that were expressed across all eight datasets (∼12,000 genes). We re-processed single-nuclei ATAC-seq datasets using the ArchR suite of tools with default parameters as outlined^129^. We extracted the `geneScoreMatrix` from the single-nuclei ATAC-seq datasets, a number that correlated with transcription of these genes that were successfully used to integrated ATAC-seq datasets with RNA-seq datasets. We used an R-package Seurat v4 to integrate the multi-omic datasets with the `SCTranform()` function to normalize gene expression with `glmGamPoi()`^130^ and employed reciprocal PCA (rPCA) method of integration `FindIntegrationAnchors()` function^131^. There was a significant difference in quality between the two rhesus macaque replicates, so they were treated as two separate datasets. We applied a guided integration that made use of known structured relatedness across the datasets. The macaque data were integrated together, followed by the human and primate datasets, and lastly the mouse dataset. After normalization and integration, every cluster had equal mixing of each dataset/species. To handle the replicate-specific batch effects within each dataset, we used the Harmony method to correct the “integrated” PCA dimensionality reduction; we also corrected for any residual batch effects not accounted for the rPCA integration^132^. We assessed the quality of the cross-species integration and sample-specific batch effect removal by creating a low-dimensionality UMAP projection plot and identified multi-species clusters using the Seurat `RunUMAP()` and `FindClusters()` functions. We visualized adequate integration and batch-correction removal. We calculated conserved marker genes for each cell type and subtype annotations. We restricted marker gene identification to snRNA-seq datasets that directly measured these transcripts. We used the `FindConservedMarker`() function. We corrected the maximum p-value for these differentially expressed genes that marked different striatal subpopulations across studies with the FDR and Bonferroni methods.

We harmonized the cell type clusters of the striatal neurons across species and used the most parsimonious cell type labels at the level of major cluster. Additionally, we annotated striatal neuron subtype levels, capturing previously appreciable biological dimensions (*e.g.*, matrix versus striosome). These labels defined a multi-species integrated annotation to standardize the reporting of single cell-omics striatal projection neurons in mammals. The labels classified SPNs in the striatum as canonical D1R^+^ and D2R^+^ SPNs as well as two transcriptomically distinct, cell types, eSPN, and IC neurons. We plotted the original cell type labels from each dataset, which represented cell type clustering at different resolutions, such as striosome versus matrix. We plotted quality control metrics to affirm none of our annotated clusters were below acceptable quality for reporting.

### Enrichment of striatal neuron cell type ATAC-seq with human genetic risk profiles

We examined whether the single-nuclei ATAC-seq profiles of striatal neuron cell types were correlated with human genetic risk variants for mental health, substance use, sleep, and other neurological traits not otherwise specified including psychosocial traits. We used the human single-nuclei ATAC-seq dataset^54^ and annotated the non-projection cell types (*e.g.*, glia, interneurons). We used `ArchR` to call reproducible peaks for D1, D2, eSPN, interneurons, astrocyte, microglia, oligodendrocyte, and oligodendrocyte precursor cells. We downloaded and processed a panel of 64 human GWAS spanning multiple domains (neurodegeneration, psychiatric disorders, substance use, psychosocial domains, sleep, and non-brain related negative control traits) (refer to Table S3 for GWAS references). We used the linkage disequilibrium score regression method (`S-LDSC`)^133^ to correlate overlap of human genome-wide association studies genetic risk variants with cell type ATAC-seq peaks as described earlier with the following exception^61^. Specifically, we used the conditional enrichment analysis version of `LDSC` to find enrichment while controlling for overlap of other striatal cell types of 200+ published features of the genome. We corrected the enrichment p-value with the Bonferroni method.

### Statistical analyses

GraphPad Prism (version 9.20, GraphPad Software, San Diego, CA) and SPSS (Version 28.01.1, IBM, Armonk, NY) were used for all statistical analyses. One-way ANOVA followed by Tukey’s multiple comparison tests were employed to analyze differences in the density of positive cells across striatal subregions. Three-way ANOVA, followed by Tukey’s post hoc comparisons, identified potential interactions between DA receptor expression, striatal subregion, and position along the rostral-caudal axis. Cohen’s D compared differences in *Foxp2* mRNA expression between striatal subpopulations.

## Supporting information

Gayden Supplementary Figures v2

## Code and Data Availability

No new sequencing experiments were performed in this study. All single-nuclei RNA-seq and single-nuclei ATAC-seq were accessed from the published accession codes or links (GSE167920, GSE147672, GSE225158, GSE139412, GSE137763, https://libd-snrnaseq-pilot.s3.us-east-2.amazonaws.com/SCE_NAc-n8_tran-etal.rda, https://data.nemoarchive.org/biccn/grant/u19_cemba/cemba/epigenome/sncell/ATACseq, http://dropviz.org). The Rscripts used to perform the cross-species and multi-dataset integration of SPNs and create the main and supplemental figures for this study are located online at https://github.com/pfenninglab/StriatumComparativeGenomics/. A copy of this code including the single-cell data files from the integration analyses with the “cell type” and spatial “subtype” labels are located at http://daphne.compbio.cs.cmu.edu/files/bnphan/StriatumComparativeGenomics/. Any intermediate files not available here can be requested from the corresponding author.

## Acknowledgments

We sincerely thank Mary Brady for assistance with figure design as well as Dr. Alan Watson, Dr. Simon Watkins and the Center for Biologic Imaging at the University of Pittsburgh for help and support with imaging and image processing. We also express gratitude to Dr. Jill Glausier for advice and guidance. We thank the authors of the single-cell studies for making their single-cell expression data and labels accessible online or via email correspondence. The species icons in Figure 6 were acquired from PhyloPic. This work was supported by NIH grants R01ES034037 (ZF), R21AG068607 (ZF), R21DA052419 (RWL, ZF), R21AA028800 (RWL, ZF), F30DA053020 (BNP), ASAP-020519 (ARP), and DP1DA046585 (ARP).

## Author Contributions

Design of the work (JG, SP, BNP, ZF); Acquisition, analysis, or interpretation of data for the work (JG, SP, BNP, CS, SAB, MCG, HAT, YD, ARP, RWL, ZF); Writing of the initial manuscript draft (JG, ZF) with other authors contributing to subsequent writing and editing.

