## Supplementary material for "Integrative multi-dimensional characterization of striatal projection neuron heterogeneity in adult brain": Gayden Supplementary Figures v2

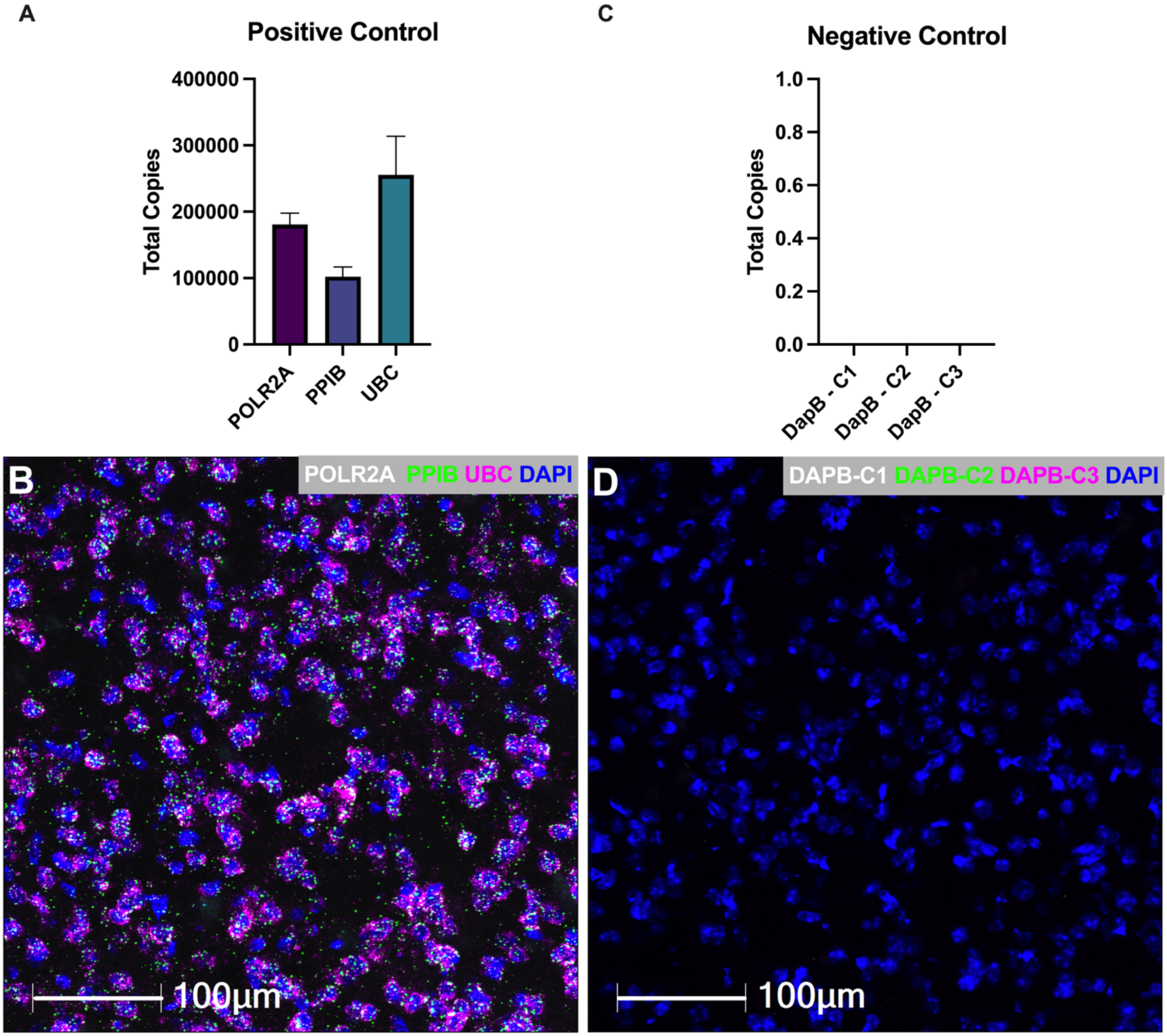


**Figure S1. Positive and negative control probes for the multiplex RNAscope assay. (A)** Quantification of the 3-plex Positive Control Probe shows a high copy number for all 3 housekeeping gene controls for each channel. **(B)** Representative RNAscope image of the positive control probe in mouse striatum, with DNA-directed RNA polymerase II subunit RPB1 (POL2RA) (shown in white), Cyclophilin B (PPIB) (shown in green), and Ubiquitin C (UBC) (shown in purple). Scale bar = 100 μm. **(C)** Quantification of the 3-plex Negative Control Probe, which targets the DapB gene in all 3 channels (DapB-C1, DapB-C2, DapB-C3), shows neither detectable staining nor background signals. **(D)** Representative RNAscope image of the negative control probe in mouse striatum. Scale bar = 100 μm.

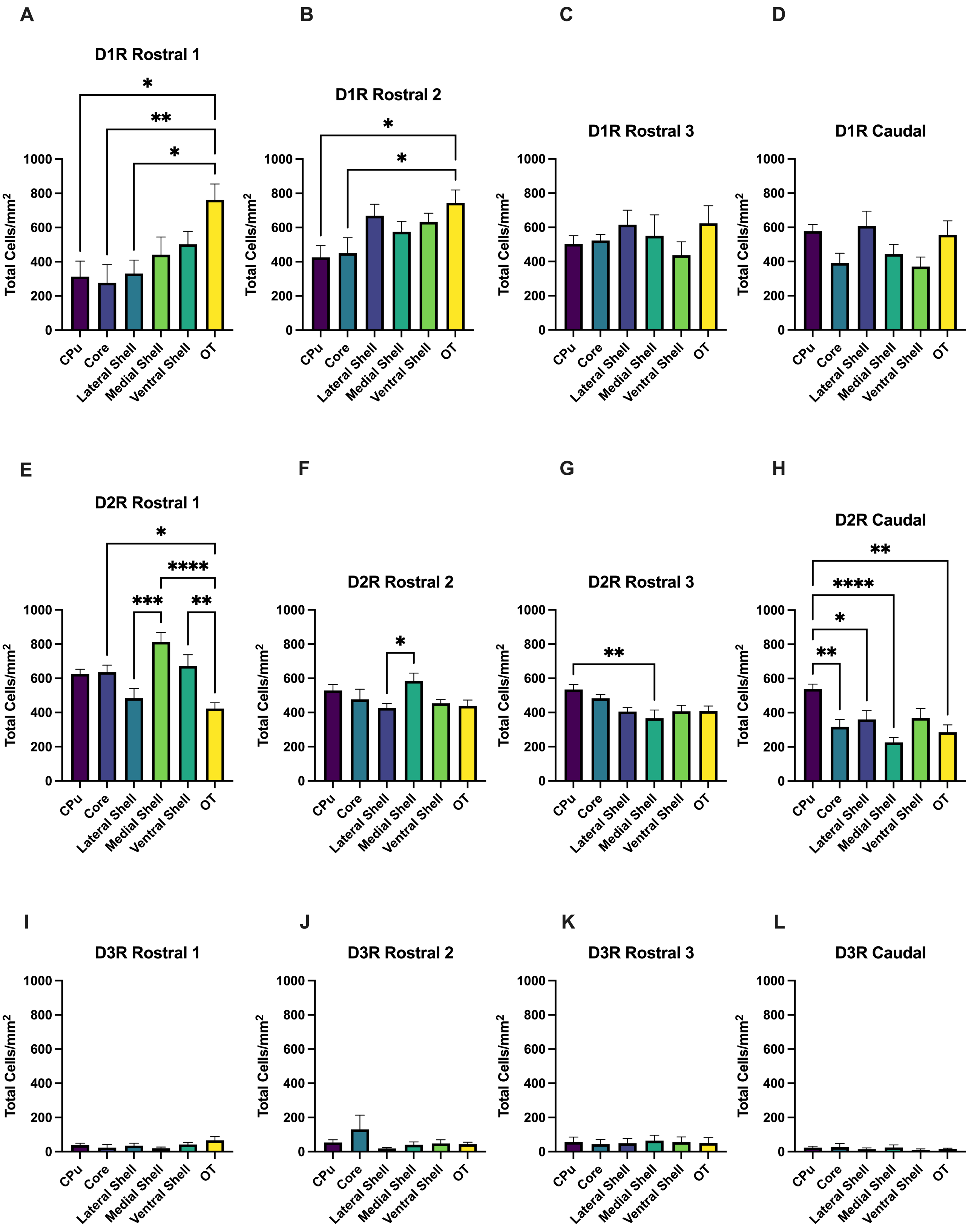
A
D1R Rostral 1
B
D1R Rostral 2
C
D1R Rostral 3
D
D1R Caudal
E
D2R Rostral 1
F
D2R Rostral 2
G
D2R Rostral 3
H
D2R Caudal
I
D3R Rostral 1
J
D3R Rostral 2
K
D3R Rostral 3
L
D3R Caudal
Total Cells/mm$^2$
0
200
400
600
800
1000
CPu
Core
Lateral Shell
Medial Shell
Ventral Shell
OT
*
**
***
****

**Figure S2. Detailed striatal subregional distribution of singly-expressing SPNs.** Division of the NAc into specific anatomic subregions (Lateral Shell, Medial Shell, Ventral Shell, and Core) enabled a more detailed analysis of the anatomic distributions of SPNs that expressed only D1R, D2R, or D3R. **(A-D)** Distribution of D1R-only SPNs shows enrichment in the OT that disappears moving caudally (*p<0.05, **p<0.01). **(E-H)** Distribution of D2R-only SPNs shows diminishing cell density in the NAc, particularly in the Medial Shell, moving caudally. **(I-L)** Distribution of D3R-only SPNs shows limited cell numbers throughout the striatum. Mean ± SEM, n=6 mice for all conditions. *p<0.05, **p<0.01, ***p<0.001, ****p<0.0001.

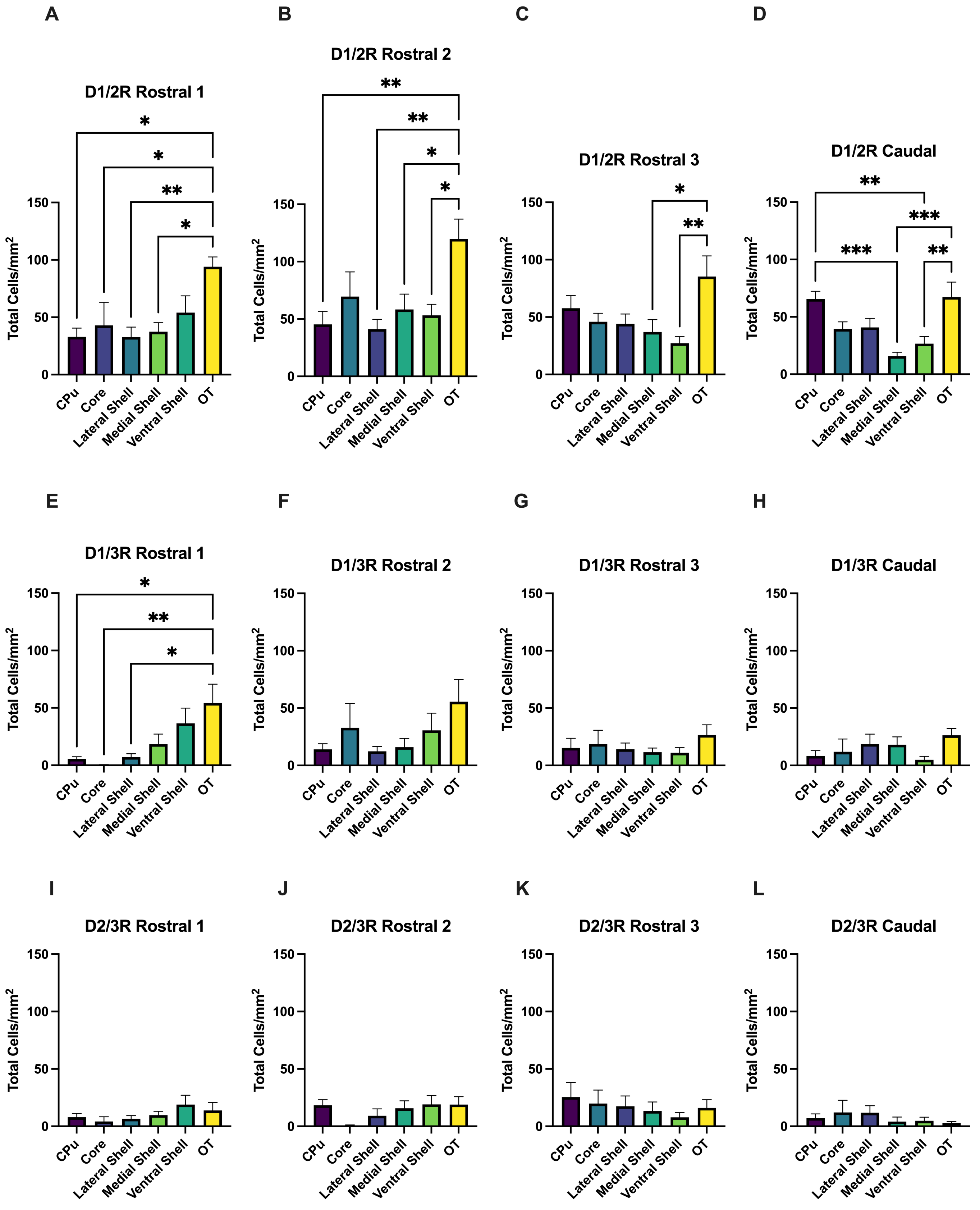
A
D1/2R Rostral 1
B
D1/2R Rostral 2
C
D1/2R Rostral 3
D
D1/2R Caudal
E
D1/3R Rostral 1
F
D1/3R Rostral 2
G
D1/3R Rostral 3
H
D1/3R Caudal
I
D2/3R Rostral 1
J
D2/3R Rostral 2
K
D2/3R Rostral 3
L
D2/3R Caudal
Total Cells/mm2
0
50
100
150
CPu
Core
Lateral Shell
Medial Shell
Ventral Shell
OT
*
**
***

**Figure S3. Detailed striatal subregional distribution of co-expressing SPNs.** Distribution of D1/2R, D1/3R, and D2/3R SPNs across the striatum across ventral-dorsal and rostral-caudal axes with the NAc divided into specific anatomic subregions. **(A-D)** Distribution of D1/2R SPNs, showing enriched expression in the olfactory tubercle (OT) throughout the rostral-caudal axis (*$p<0.05$, **$p<0.01$, ***$p<0.001$). **(E-H)** Distribution of D1/3R SPNs reveals a strong enrichment in the OT in the most rostral section **(E)** that is not evident in more caudal sections (*$p<0.05$, **$p<0.01$). **(I-L)** Distribution of D2/3R SPNs shows low levels of cells even distributed throughout the striatum. Mean ± SEM, n=6 mice for all conditions. *$p<0.05$, **$p<0.01$, ***$p<0.001$.

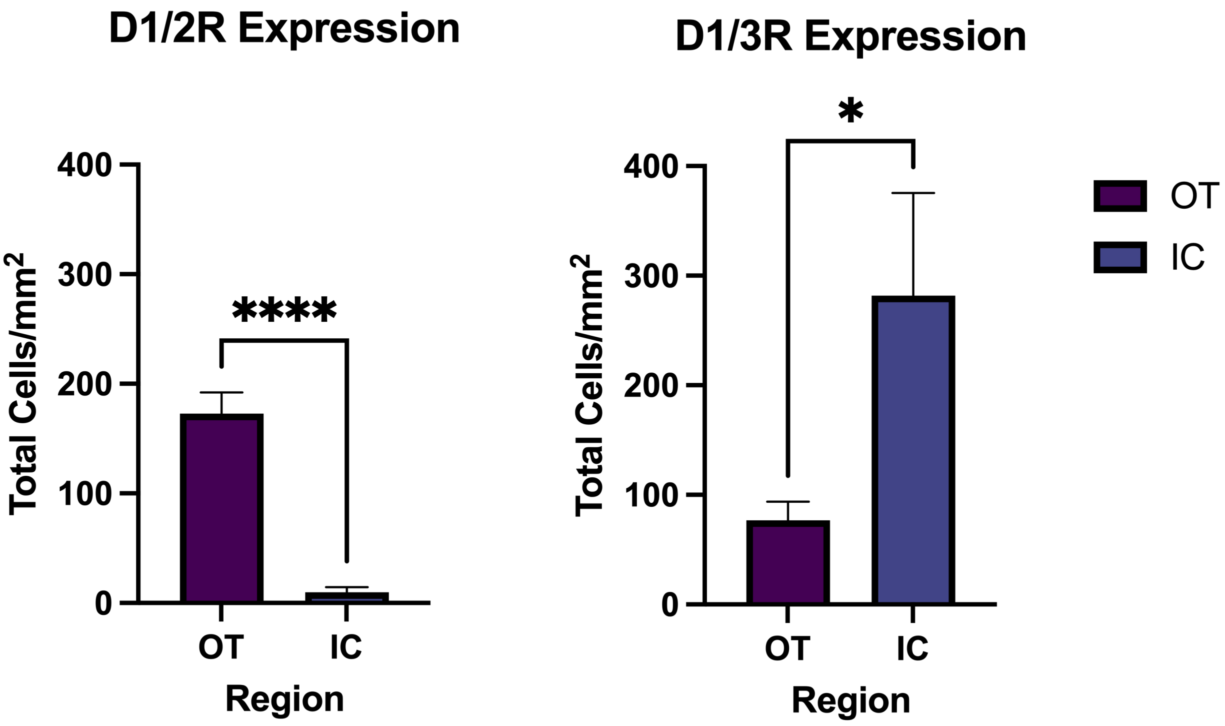


**Figure S4. Spatial distribution of D1/2R and D1/3R co-expressing SPNs within the ventralmost striatum.** Quantification of D1/2R and D1/3R co-expressing SPNs in most ventral subregion of the striatum including the olfactory tubercle (OT) and Islands of Calleja (IC). D1/2R SPNs were predominantly expressed in the OT as a whole compared to the IC (****p<0.0001). D1/3R SPNs were significantly enriched in the IC versus the OT (*p=0.0237). Mean ± SEM, n=6 mice for all conditions. *p<0.05, **p<0.01, ***p<0.001, ****p<0.0001.

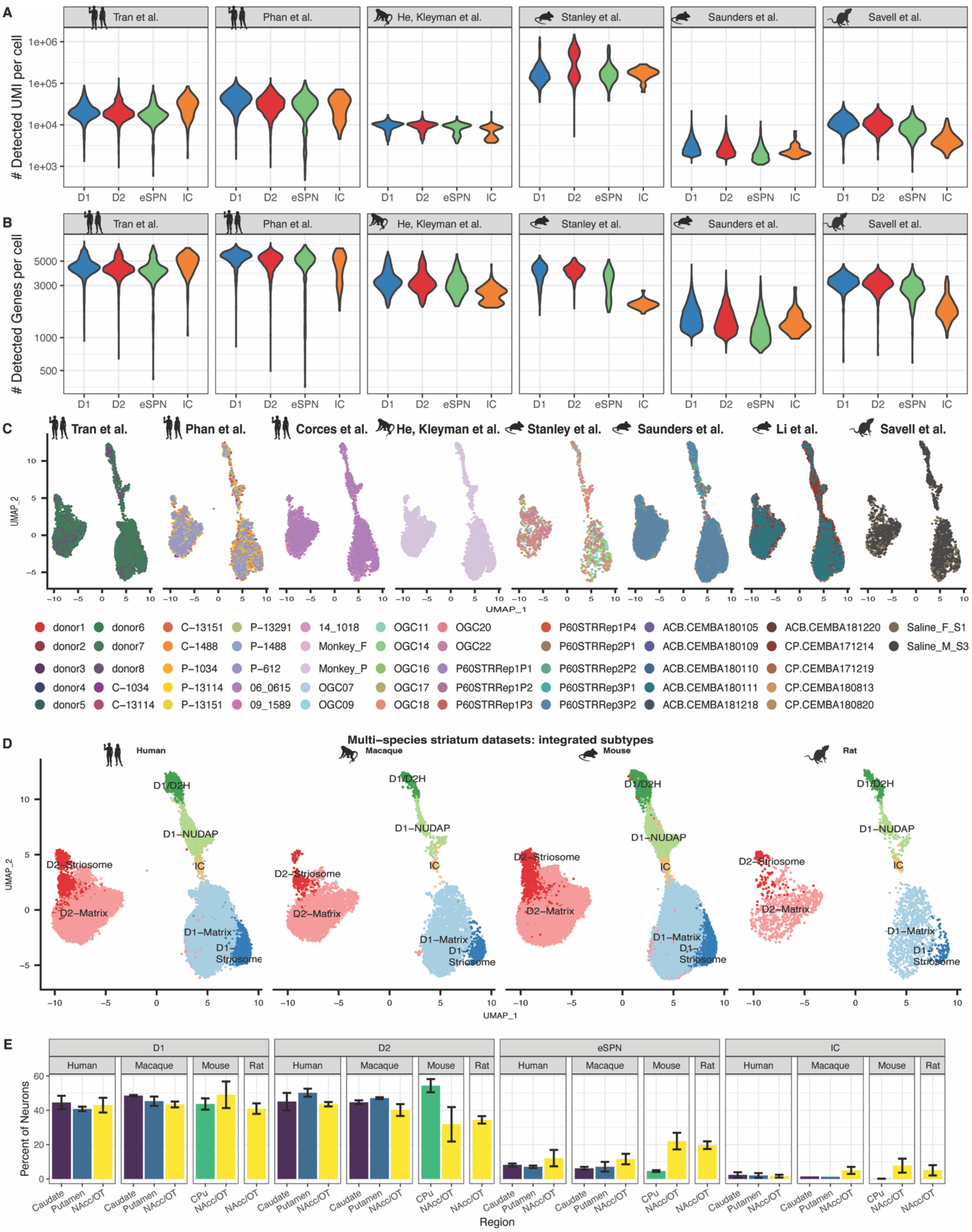
A
Tran et al.
Phan et al.
He, Kleyman et al.
Stanley et al.
Saunders et al.
Savell et al.
### Detected UMI per cell
1e+06
1e+05
1e+04
1e+03
D1 D2 eSPN IC
B
### Detected Genes per cell
5000
3000
1000
500
C
Tran et al.
Phan et al.
Corces et al.
He, Kleyman et al.
Stanley et al.
Saunders et al.
Li et al.
Savell et al.
UMAP_2
UMAP_1
donor1 donor2 donor3 donor4 donor5 donor6 donor7 donor8
C−13151 C−1488 P−1034 P−13114 P−13151 C−1034 C−13114
P−13291 P−1488 P−612 06_0615 09_1589
14_1018 Monkey_F Monkey_P
OGC07 OGC09 OGC11 OGC14 OGC16 OGC17 OGC18 OGC20 OGC22
P60STRRep1P1 P60STRRep1P2 P60STRRep1P3 P60STRRep1P4 P60STRRep2P1 P60STRRep2P2 P60STRRep3P1 P60STRRep3P2
ACB.CEMBA180105 ACB.CEMBA180109 ACB.CEMBA180110 ACB.CEMBA180111 ACB.CEMBA181218 ACB.CEMBA181220
CP.CEMBA171214 CP.CEMBA171219 CP.CEMBA180813 CP.CEMBA180820
Saline_F_S1 Saline_M_S3
D
Multi−species striatum datasets: integrated subtypes
Human
Macaque
Mouse
Rat
D1/D2H
D1−NUDAP
IC
D2−Striosome
D2−Matrix
D1−Matrix
D1−Striosome
UMAP_2
UMAP_1
E
D1
D2
eSPN
IC
Human Macaque Mouse Rat
Percent of Neurons
60
40
20
0
Caudate Putamen NAcc/OT CPu
Region

**Figure S5. Sample-wise quality control metrics and integration. (A)** Violin plot of unique mRNAs detected per cell stratified by cell type and single-cell RNA-seq dataset. **(B)** Violin plot of genes detected per cell stratified by cell type and single-cell RNA-seq dataset. **(C)** UMAP projection plot of integrated single-cell RNA-seq and ATAC-seq datasets colored by biological replicates stratified by dataset. **(D)** UMAP projection plot of integrated single-cell RNA-seq and ATAC-seq datasets colored by striatal projection neuron subtypes stratified by species. **(E)** Proportion of transcriptomically defined neurons across the striatum. Percent of each neuron subtype across species and striatal subregions; data is represented as mean ± SEM.

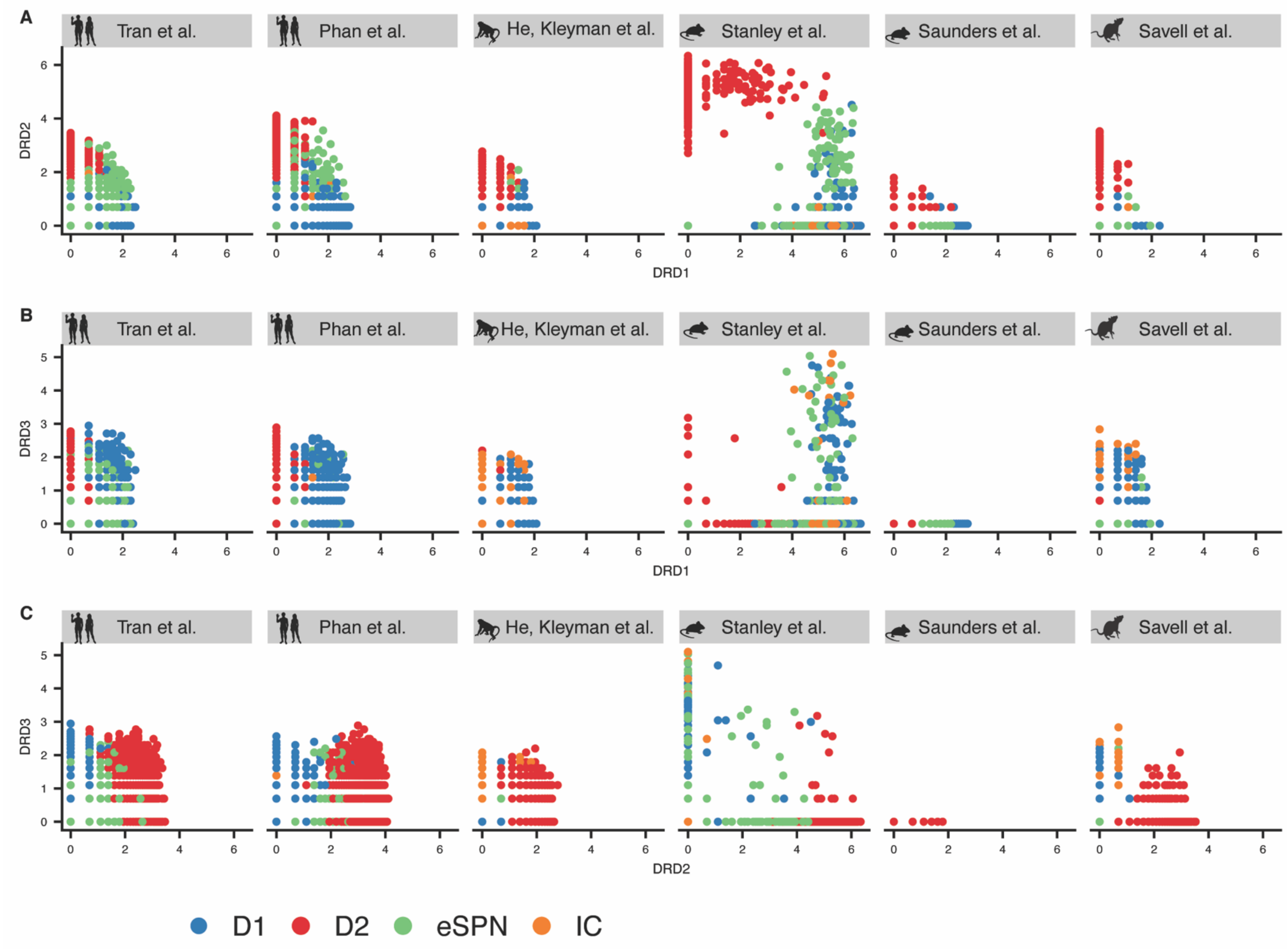


**Figure S6. Co-expression of dopamine receptors across single-nuclei RNA-seq datasets.** Scatterplot of dopamine receptor expression within single cells across all six analyzed studies colored by consensus cell type labels. **(A)** Plot of *DRD1* and *DRD2*, **(B)** Plot of *DRD1* and *DRD3*, and **(C)** Plot of *DRD2* and *DRD3*.
