## Supplementary material for "Integrative multi-dimensional characterization of striatal projection neuron heterogeneity in adult brain": Table S1

| Rostral 1 | CPu | Core | Shell | OT |
|---|---|---|---|---|
| **D1/D2+ out of total D1+ cells** | 10.6 ± 1.5 | 8.2 ± 2.5 | 9.3 ± 1.0 | 14.8 ± 1.4 |
| **D1/D2+ out of total D2+ cells** | 6.7 ± 1.9 | 6.0 ± 2.2 | 6.5 ± 0.8 | 21.2 ± 2.2 |
| **D1/D3 out of D1+ cells** | 2.2± 0.9 | 0.0 ± 0.0 | 7.0 ± 1.9 | 7.4 ± 2.3 |
| **D2/D3+ out of D2+ cells** | 1.2 ± 0.5 | 0.0 ± 0.0 | 1.9 ± 0.5 | 5.0 ± 2.1 |

**Table 1.1:** Percentage of striatal medium spiny neurons that co-express dopamine receptors in rostral 1 section (Bregma-1.61mm)

| Rostral 2 | CPu | Core | Shell | OT |
|---|---|---|---|---|
| **D1/D2+ out of D1+ cells** | 11.6 ± 1.8 | 10.8 ± 2.7 | 8.2 ± 0.7 | 15.5 ± 1.5 |
| **D1/D2+ out of D2+ cells** | 8.0 ± 1.7 | 10.4 ± 2.9 | 10.2 ± 1.0 | 28.1 ± 3.9 |
| **D1/D3 out of D1+ cells** | 3.3 ± 0.9 | 0.1 ± 0.1 | 1.9 ± 0.4 | 7.1 ± 1.8 |
| **D2/D3+ out of D2+ cells** | 3.8 ± 1.1 | 0.0 ± 0.0 | 1.8 ± 0.5 | 2.7 ± 0.8 |

**Table 1.2:** Percentage of striatal medium spiny neurons that co-express dopamine receptors in rostral 2 section (Bregma -1.33mm)

| Rostral 3 | CPu | Core | Shell | OT |
|---|---|---|---|---|
| **D1/D2+ out of D1+ cells** | 10.6 ± 1.2 | 8.8 ± 1.1 | 6.4 ± 0.6 | 12.9 ± 1.3 |
| **D1/D2+ out of D2+ cells** | 10.3 ± 1.7 | 9.5 ± 1.4 | 8.4 ± 1.0 | 19.9 ± 3.9 |
| **D1/D3 out of D1+ cells** | 0.8 ± 0.4 | 0.0 ± 0.0 | 1.2 ± 0.3 | 3.6 ± 1.1 |
| **D2/D3+ out of D2+ cells** | 1.6± 0.9 | 0.2 ± 0.2 | 1.2 ± 0.5 | 2.6 ± 1.1 |

**Table 1.3:** Percentage of striatal medium spiny neurons that co-express dopamine receptors in rostral 3 section (Bregma -1.15mm)

| Caudal | CPu | Core | Shell | OT |
|---|---|---|---|---|
| **D1/D2+ out of D1+ cells** | 11.3 ± 0.8 | 10.9 ± 2.3 | 5.7 ± 0.8 | 11.9 ± 2.2 |
| **D1/D2+ out of D2+ cells** | 11.9 ± 0.9 | 12.8 ± 2.0 | 8.9 ± 1.1 | 26.0 ± 5.7 |
| **D1/D3 out of D1+ cells** | 0.8 ± 0.3 | 0.1 ± 0.1 | 2.1 ± 0.6 | 3.9 ± 0.8 |
| **D2/D3+ out of D2+ cells** | 0.4 ± 0.2 | 0.0± 0.0 | 0.0± 0.0 | 0.2 ± 0.2 |

**Table 1.4:** Percentage of striatal medium spiny neurons that co-express dopamine receptors in the caudal section (Bregma +0.91mm)

**Table S1. Percentage of dopamine $D_1$, $D_2$, and $D_3$ receptors co-expressed in $D_1R^+$ and $D_2R^+$ SPNs across ventral-dorsal and rostral-caudal axes.**
